# Metabolomics of bacterial-fungal pairwise interactions reveal conserved molecular mechanisms

**DOI:** 10.1101/2023.03.13.532449

**Authors:** Gordon T. Luu, Jessica C. Little, Emily C. Pierce, Manon Morin, Celine A. Ertekin, Benjamin E. Wolfe, Oliver Baars, Rachel J. Dutton, Laura M. Sanchez

**Affiliations:** Department of Chemistry and Biochemistry, University of California Santa Cruz, Santa Cruz, California, 95064; Department of Pharmaceutical Sciences, University of Illinois at Chicago, Chicago, Illinois, 60612; Division of Biological Sciences, University of California San Diego, La Jolla, California, 92093; Center for Microbiome Innovation, Jacobs School of Engineering, University of California San Diego, La Jolla, 92093; Department of Biology, Tufts University, Medford, Massachusetts, 02155; Tufts University Sensory and Science Center, Medford Massachusetts, 02155; Department of Entomology and Plant Pathology, North Carolina State University, Raleigh, North Carolina, 27607

## Abstract

Bacterial-fungal interactions (BFIs) can shape the structure of microbial communities, but the small molecules mediating these BFIs are often understudied. We explored various optimization steps for our microbial culture and chemical extraction protocols for bacterial-fungal co-cultures, and liquid chromatography-tandem mass spectrometry (LC-MS/MS) revealed that metabolomic profiles are mainly comprised of fungi derived features, indicating that fungi are the key contributors to small molecule mediated BFIs. LC-inductively coupled plasma MS (LC-ICP-MS) and MS/MS based dereplication using database searching revealed the presence of several known fungal specialized metabolites and structurally related analogues in these extracts, including siderophores such as desferrichrome, desferricoprogen, and palmitoylcoprogen. Among these analogues, a novel putative coprogen analogue possessing a terminal carboxylic acid motif was identified from *Scopulariopsis* spp. JB370, a common cheese rind fungus, and its structure was elucidated via MS/MS fragmentation. Based on these findings, filamentous fungal species appear to be capable of producing multiple siderophores with potentially different biological roles (i.e. various affinities for different forms of iron). These findings highlight that fungal species are important contributors to microbiomes via their production of abundant specialized metabolites and their role in complex communities should continue to be a priority.

## Introduction

One of the major driving factors behind complex microbial communities are the interactions between bacteria and fungi^1^. Both domains are commonly found to co-occur in and on animals, soil, and plants; yet the specialized metabolites that mediate interactions between bacteria and fungi are understudied, especially in the case of commensal fungi^2,3^. Studies suggest that bacterial-fungal interactions (BFIs) are mediated by small molecules, and these interactions may impact how microbiomes form and function^2–6^.

Metabolites represent the net result of gene regulation and can serve as a functional readout for biosynthesis from their cognate proteins^7,8^. The observed effects of metabolites can often be categorized as direct or indirect^9^. Direct effects are readily observable and have inspired the field of antimicrobial drug discovery, whereas indirect effects typically involve factors such as a change in environmental pH, altered nutrient availability, or stimulation of a host immune response, and are often more challenging to measure and assess. There are many described mutualistic relationships between bacteria and fungi,^11^ and in many cases these relationships involve the exchange of metabolites. Fungi respond to bacterial metabolites with secretion of their own metabolites and vice versa^12^. The result is a dynamic chemical interplay between bacteria and fungi^13^. In many cases, the presence of naturally occurring microbial cohabitants is imperative for the production of compounds encoded by cryptic or quiet biosynthetic gene clusters (BGCs) that are otherwise undetectable in monoculture^14–17^.

The present study expands on our previous work using a model microbiome - cheese rinds - to systematically evaluate the effects of metabolites exchanged between bacteria and fungi. The cheese rind microbiome has been shown to be a simplified model for the study of complex microbial interactions based on relatively reduced complexity and reproducibility^18^. High correlation between bacterial and fungal genera found on cheese rinds cannot be explained by environmental factors alone, which suggests that mechanisms beyond simple pH and salt content changes drive the predictable co-occurrence of genera^19^. Specialized metabolites are likely central to BFIs on cheese rinds^20^.

Previously, high-throughput screening of pairwise bacterial-fungal combinations was used to select strains of interest based on their phenotypic properties in co-culture such as pigment production, altered nutrient requirements, delayed or promoted growth, and inhibition. Six filamentous fungal strains (*Penicillium solitum* #12, *Penicillium camembertii* SAM3, *Penicillium atramentosum* RS17, *Scopulariopsis* sp. JB370, *Scopulariopsis* sp. 165-5, and *Fusarium domesticum* 554A) and two yeast strains (*Diutina catenulata* 135E and *Debaromyces hansenii* 135B) were grown in pairwise co-culture with one of two bacterial growth partners: *Pseudomonas psychrophila* JB418, which is a member of the cheese rind microbiome, and *Escherichia coli* K12. Random barcode transposon-site sequencing (RB-TnSeq) revealed that fungi impacted the fitness of bacterial mutants compared to those of bacterial monocultures through conserved mechanisms including changes of iron and biotin availability and the presence of fungi-derived antibiotics^21^. In the present study, we use these same bacterial-fungal co-cultures to systematically study the metabolome using liquid chromatography tandem mass spectrometry (LC-MS/MS) and found that fungi are prominent producers of specialized metabolites. We have also putatively identified a unique siderophore produced by *Scopulariopsis* sp. JB370. Finally, we have looked at the effects that different culture and extraction methods have on mass spectrometry-based untargeted metabolomics experiments, which can inform on how to handle diverse microbes in complex interkingdom co-cultures.

## Materials and Methods

### Microbial Culturing for Specialized Metabolite Extraction

All bacterial cultures were grown overnight in brain heart infusion (Bacto; BHI) liquid media at room temperature. Liquid cultures were normalized to an optical density (OD_600_) of 0.1 and bacterial cultures were diluted 10^−1^ for further experiments. Fungal cultures were grown on plate count agar milk salt (1 g/L whole milk powder, 1 g/L dextrose, 2.5 g/L yeast extract, 5 g/L tryptone, 10 g/L sodium chloride, 15 g/L agar; PCAMS)^22^. Plates were kept at room temperature and spores were harvested at 7 days (or until sporulation was observed) of growth for subsequent experiments. Spores harvested from fungi were put into 1X PBS and normalized to OD_600_ 0.1 for further experiments.

For all cultures, three biological replicates of each condition were plated and extracted from solid agar. For extraction of solid agar plates, 5 µL of working cultures of both bacteria and fungi were spotted onto 10% cheese curd agar medium (100 g/L freeze dried cheese curd, 5 g/L xanthum gum, 30 g/L sodium chloride, 17 g/L agar; CCA) adjusted to pH 7. For co-cultures, bacteria and fungi were streaked as t-spots with two different co-culture spots per plate varying the orientation of bacteria and fungi (**Figure S1**). For monocultures, bacteria and fungi were streaked onto plates with two spots per plate. After at least 7 days of growth, agar was removed from the petri plate and placed into 50 mL conical tubes. Acetonitrile (10 mL) was added to each tube and all were sonicated for 30 min. All conical tubes were centrifuged at 4000 rpm, 4 °C for 15 min and the liquid supernatant was removed from the solid agar pieces and put into 15 mL conical tubes. The 15 mL conical tubes containing liquid were then centrifuged at 4000 rpm, 4 °C for 15 min and liquid was again removed and stored. These liquid extractions were then dried *in vacuo*. Dried extracts were resuspended in 90:10 methanol:DI water and centrifuged to pellet insoluble material. These extracts were dried *in vacuo* again, weighed and resuspended in 90:10 methanol:DI water to obtain 1 mg/mL solutions and analyzed via LC-ICP-MS and LC-MS/MS.

Subsequent monocultures and co-cultures used to test the inclusion of Actinobacteria in pairwise co-cultures and the performance of plug extractions were subjected to the same culture and (full plate) extraction conditions. Additionally, a modified plug extraction method was used on a set of bacterial and fungal monocultures as follows. After at least 7 days of growth, a plug was removed by stamping the agar with a 50 mL conical tube to yield a 30 mm plug (as opposed to removing the entire plate) and placed into 50 mL conical tubes. 10 mL acetonitrile was added to each tube and all were sonicated for 30 min. All conical tubes were centrifuged at 4000 rpm, 4 °C for 15 min and the liquid supernatant was removed from the solid agar pieces and dried *in vacuo*. Samples were resuspended in 50:50 methanol:DI water and diluted to a concentration of 1 mg/mL solutions, which were then transferred to LC vials and analyzed via LC-MS/MS.

Cultures performed on nitrocellulose membranes were grown on PCAMS and subject to the same culture conditions described above, with the exception that one set of samples were inoculated on top of a nitrocellulose membrane that was activated and placed on top of all agar surfaces. After at least 7 days of growth, cultures without a nitrocellulose membrane were extracted as described above using the full plate extraction (albeit from 60 mm plates). Cultures with a nitrocellulose membrane had the microbial biomass completely removed and were extracted as described above using the full plate extraction. All extracts were dried *in vacuo* and resuspended in 50:50 methanol:DI water and diluted to a concentration of 1 mg/mL and analyzed via LC-MS/MS.

### Actinobacteria Strain Prioritization Using IDBac

*Glutamicibacter arilaitensis* JB182, 12 *Brevibacterium* strains, 3 *Brachybacterium* strains, *Hafnia alvei* JB232, *P. psychrophila* JB418, and *E. coli* K12 were grown on PCAMS at room temperature. Bacterial stocks were harvested to create a 12.5% glycerol stock and stored at -70 °C for subsequent experiments. Eight biological replicates for each strain were then inoculated (5 µL) onto 10% CCA, incubated at rt, and processed as described below.

### Chrome Azurol S (CAS) Assay for Detection of Siderophores

For preparation of the CAS reagent, polypropylene pipette tips and conical tubes were used in preparation of reagents to avoid contact with glass. A 2 mM CAS stock solution was prepared by adding CAS to DI water. A 1 mM FeCl_3_ stock solution was prepared by adding FeCl_3_ 6 H_2_O to 10 mM HCl. Piperazine buffer was prepared by dissolving anhydrous piperazine in water and adding 5 M HCl until the pH reached 5.6. A 5-sulfosalicylic acid solution (0.2 M) was prepared by adding 5-sulfosalicylic acid to DI water. To prepare 10 mL of CAS reagent, 0.00192 g of hexadecyltrimethylammonium bromide (HDTMA) was added to 5 mL of MilliQ water in a 50 mL conical tube; while stirring, 750 µL of CAS solution, 150 µL of 1 mM FeCl_3_, 3 mL of piperazine solution, and 200 µL 5-sulfosalicylic acid solution were added, the total volume of the solution was brought up to 10 mL by the addition of DI water. The resulting CAS assay reagent and solution ingredients were stored in the dark at 4 °C.

To perform the CAS assay, monoculture extracts were dissolved in 90:10 water:methanol and centrifuged to pellet any solid material. Dissolved material were transferred and weighed to achieve a 1 mg/mL solution. Solutions (1 mg/mL) were serially diluted. Deferoxamine-mesylate standard (1 M solution,Sigma-Aldrich) was used as a positive control and serially diluted to obtain decreasing concentrations. Sample and CAS assay reagents were mixed by pipetting in a 1:1 volume ratio (100 µL sample:100 µL reagent) and incubated at room temperature for 120 min. The optical density at 630 nm (OD_630_) was measured in a 96-well plate at the end of 120 min.

### LC-MS Data Collection for Quantification of Metabolites from Bacterial-Fungal Co-Cultures

High resolution LC-MS data for individual biological replicates of bacterial-fungal monocultures and pairwise co-cultures was collected on a Bruker compact qTOF in positive mode with a detection window set from 100 - 2000 Da on an Agilent 2.1 × 50 mm C18 Poroshell UPLC column with a flow rate of 0.5 mL/min. A gradient of using MilliQ water with 0.1% formic acid (solvent A) and acetonitrile with 0.1% formic acid (solvent B) was used with an initial 1 minute hold at 10% B, 10 - 100% B over 12 minutes, and a 1 minute hold at 100% B, followed by a 1 minute equilibration. Agilent ESI TOF mix calibrant was injected via divert valve from minute 0.04 to 0.30 during the LC run. For each sample, 10 µL of a 1 mg/mL solution was injected. This dataset will be referred to as Dataset 1 from here on out.

### LC-MS(/MS) Data Collection for Relative Quantification and Annotation of Metabolites to Evaluate Actinobacteria Prioritization, Actinobacteria Specialized Metabolite Production, Plug Extractions, Nitrocellulose Cultures

High resolution LC-MS/MS data were collected on a Bruker timsTOF fleX in positive mode. The detection window was set from 100 to 2000 Da. An Agilent Poroshell 120 Å 2.1 × 50 mm UPLC column was used with a flow rate of 0.5 mL/min where solvent A was MilliQ H_2_O w/ 0.1% formic acid and solvent B was acetonitrile w/ 0.1% formic acid. Data were acquired in Compass Hystar 6.0 and otofControl 6.2. The following gradient was used: hold 5% B from 0 to 2 min, 5 - 100% B from 2 to 12 min, and hold 100% B from 12 to 14 min, followed by a 1 min equilibration time. For each sample, 2 μL of a 1 mg/mL solution was injected in partial loop mode with a loop volume set to 50 μL. The ESI conditions were set with the capillary voltage at 4.5 kV. For MS/MS, dynamic exclusion and the top nine precursor ions from each MS^1^ scan were subject to collision energies scaled according to mass and charge state for a total of nine data dependent MS/MS events per MS^1^ via Auto MS/MS mode. The datasets generated to prioritize Actinobacteria strains, evaluate Actinobacteria growth partners, test the performance of plug extractions, and test the performance of colony removal with nitrocellulose membranes will be referred to as Dataset 2, 3, 4, and 5, respectively, from here on out.

### LC-ICP-MS Data Collection

LC separation was performed on a Thermo Dionex Ultimate 3000 UPLC system using an ACE excel 1.7 super C18 2.1 × 100 mm UPLC column with a flow rate of 0.35 mL/min. The column outflow was split using a flow-splitter (Analytical Scientific Instruments, 600-PO10-06) to introduce 0.1 mL min1 into the ICP-MS with a zero dead volume PFA micro-nebulizer (Elemental Scientific). Solvent A was MilliQ water with 5 mM ammonium formate adjusted to pH 6.5 using ammonium hydroxide. Solvent B was 95:5 ACN:MilliQ water with 5 mM ammonium formate adjusted to pH 6.5 using ammonium hydroxide. Solvent was held at 0% B for 2 minutes and then a gradient of 0 - 100% B over 15 minutes was run and brought back down to 0% B at 15.2 minutes. For each sample, 20 uL of a 1 mg/mL solution was injected.

The LC flow was connected to a computer-controlled Thermo iCAP RQ ICPMS controlled by QTEGRA and operating in LC mode for ICP-MS data collection. Additional oxygen gas flow to the inlet was set to 4.50%.. The instrument was operated in KED mode with He as a collision gas. The isotopes chosen for analysis were ^48^Ti, ^55^Mn, ^56^Fe, ^57^Fe, ^59^Co, ^60^Ni, ^63^Cu, ^64^Zn, ^65^Cu, ^66^Zn, and ^95^Mo, because these transition metals have been found chelated to microbial ionophores. This dataset will be referred to as Dataset 6 from here on out.

### LC-MS/MS Data Collection for Correlation to LC-ICP-MS Data

High resolution LC-MS/MS data for pooled biological replicates (N=3) was collected on a Bruker compact qTOF in positive mode with the detection window set from 100 - 2000 Da on an ACE excel 1.7 super C18 2.1 × 100 mm UPLC column with a flow rate of 0.35 mL/min. Agilent ESI TOF mix calibrant was injected via divert valve from minute 0.04 to minute 0.30 during the LC run. Solvent A was MilliQ water with 5 mM ammonium formate adjusted to pH 6.5 using ammonium hydroxide. Solvent B was 95.5 ACN:MilliQ water with 5 mM ammonium formate adjusted to pH 6.5 using ammonium hydroxide. Solvent was held at 0% B for 2 minutes and then a gradient of 0 - 100% B over 15 minutes was run and brought back down to 0 % at 15.2 minutes. For each sample, 20 uL of a 1 mg/mL solution was injected. The ESI conditions were set with a capillary voltage at 4.5 kV. For MS/MS, dynamic exclusion was used, and the top nine precursor ions from each MS^1^ scan were subjected to collision energies scaled according to mass and charge state for a total of nine data dependent MS/MS events per MS^1^. This Dataset will be referred to as Dataset 7 from here on out.

### IDBac Data Collection via MALDI-TOF MS for Actinobacteria Strain Prioritization

After at least 7 days of growth, a chemical extraction protocol was performed on biological replicates (N=4) of each strain as previously reported^23^. Data acquisition settings were based off of those previously developed by Clark *et al*.^24^.

For analysis of intact protein profiles via MALDI-TOF MS, the supernatant was co-crystallized 1:1 with 20 mg/mL sinapinic acid in 70:30 ACN:DI water with 0.1% TFA and 1.5 µL of each sample was spotted onto a MALDI ground steel plate. Measurements were performed using a Bruker Microflex LT mass spectrometer equipped with a nitrogen laser. Protein profiles were collected in positive linear mode (600 shots; Frequency: 60 Hz) with a mass range of 2000 - 20000 Da. Protein spectra were corrected using Protein Standard I (Bruker Daltonics) as an external calibrant. Automated data acquisition was performed using flexControl v3.4 (Bruker Daltonics) and flexAnalysis v3.4. Spectra were automatically evaluated during acquisition to determine whether a spectrum was of high enough quality to retain and add to the sum of the sample acquisition.

For analysis of small molecule profiles via MALDI-TOF MS, the supernatant was co-crystallized 1:1 with a supersaturated solution of 50:50 α-cyano-4-hydroxycinnamic acid:2,5-dihydrobenzoic acid (CHCA:DHB) in 78:22 ACN:DI water with 0.1% TFA and 1.5 µL of each sample was spotted onto a MALDI ground steel plate. Measurements were performed using a Bruker timsTOF fleX mass spectrometer equipped with a smartbeam 3D™ laser (355 nm). Small molecule profiles were collected in positive reflectron mode (5000 shots; Frequency: 1000 Hz) with a mass range of 100 - 2000 Da. Small molecule spectra were corrected using Peptide Standard I (Bruker Daltonics) as an external calibrant. Automated data acquisition was performed using timsControl v2.0 (Bruker Daltonics). This dataset will be referred to as Dataset 8 from here on out.

### Data Availability

Full scan MS^1^ data for Dataset 1 can be found under MassIVE accession number MSV000085046. MS/MS data from pooled biological replicates from Dataset 1 can be found under MassIVE accession number MSV000085070 and MS/MS data from individual biological replicates can be found under accession number MSV000085007. Dataset 2 can be found under MassIVE accession number MSV000091358. Dataset 3, 4, and 5 can be found under MassIVE accession number MSV000091360. Dataset 6 can be found under MassIVE accession number MSV000091384. Dataset 7 can be found under MassIVE accession number MSV000091385. Dataset 8 can be found under MassIVE accession number MSV000089824 (protein profiles) and MSV000091361 (small molecule profiles). The GNPS feature based molecular networking (FBMN) job from which **Figure 6** is derived can be found at https://gnps.ucsd.edu/ProteoSAFe/status.jsp?task=345b589dcf84403489b04458cc94c40a.

## Results and Discussion

In order to assess the microbial strains for specialized metabolite production, mass spectrometry was used to establish baseline metabolomic profiles of all 10 strains of bacteria and fungi followed by the measurement of metabolic changes in response to the presence of a growth partner. All fungal strains were co-cultured with one of the two bacterial partners and the resulting metabolomic profiles were evaluated based on the presence or absence of features detected via LC-MS/MS and differential expression of these features based on changes in culture community composition.

### Fungal Metabolites are Prominent, Significant Features in Metabolome Profiles

To determine (dis)similarity in the overall metabolomic profiles, the data from Dataset 1 were processed using MZmine2 to create feature lists, containing an *m/z* and retention time pairing, found across all samples^25^. These feature lists were then considered individual metabolomic profiles for the given growth conditions tested (monoculture or co-culture). Hierarchical clustering of the samples was performed to compare global aspects of these profiles using the MetaboAnalyst platform^26^. To visualize how features contributed to this clustering, the biological replicates of each condition were averaged and displayed according to hierarchical clustering of samples and features in a heatmap (**Figure 1**). The heatmap represents feature intensities across all samples and considers only the most significant features (top 1000) as determined by the analysis of variance test (ANOVA).

**Figure 1:**
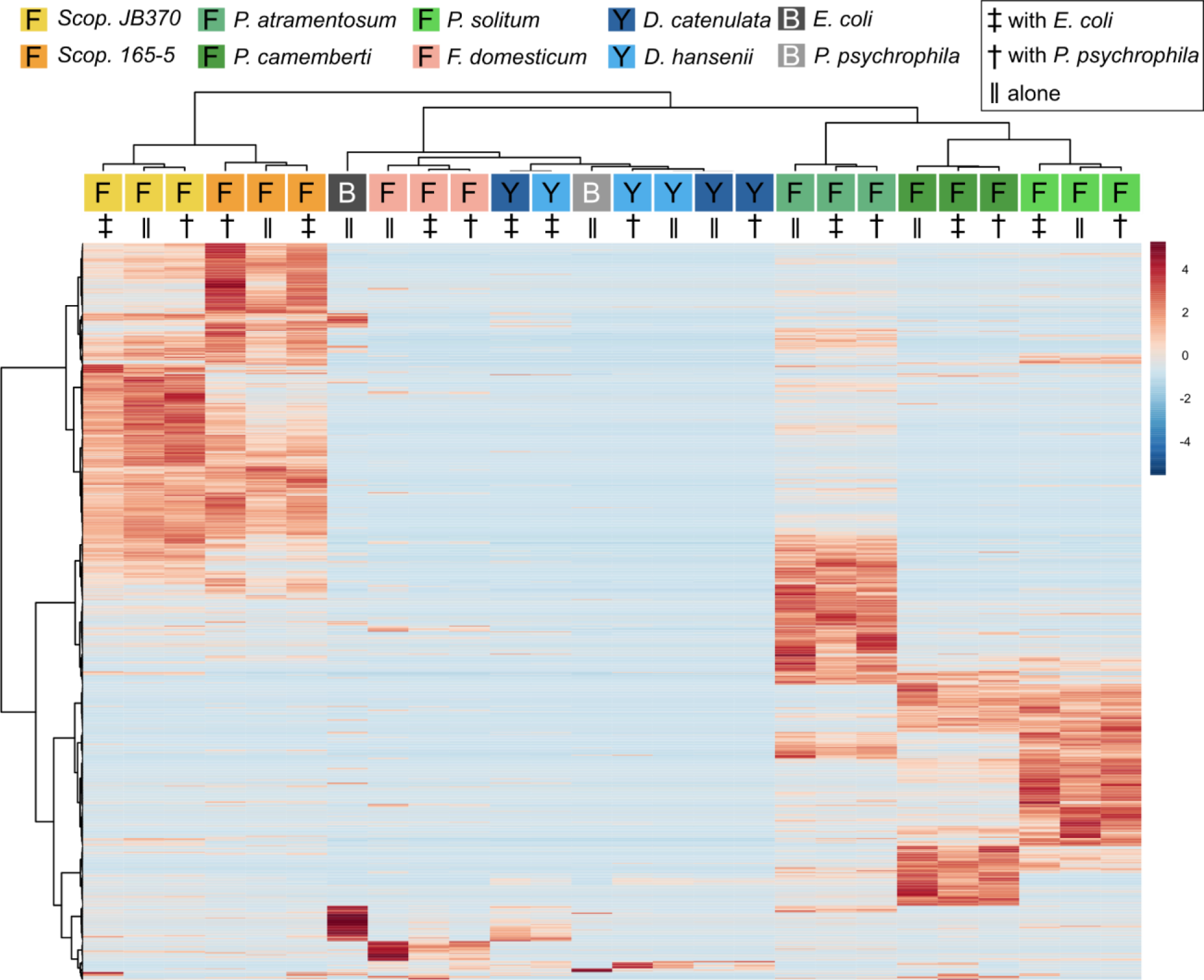
Hierarchical clustering of features and samples is displayed in this heatmap. Cultures grown here were grown with exposure to light. The top 1000 most significant features as determined by ANOVA are plotted by intensity in the sample, and only group averages are shown here with a group referring to the average of all biological replicates across one sample condition. Sample and feature clustering was performed using the Ward algorithm based on Euclidean distance and samples are colored according to fungal genera. Species were also color coded based on their genera (i.e. *Scopulariopsis* are different shades of yellow/orange, *Penicillium* are different shades of green, etc.); furthermore, species denoted with an “F” are indicative of filamentous fungi, species denoted with a “Y” are a yeasts, and species denoted with a “B” are bacteria. Symbols are under the colored terminus of the fungal dendrogram to indicate whether fungi were grown alone or with a bacterial growth partner.

Overall, with the exception of the yeast species, metabolomes cluster together based on species of fungi present rather than whether they were cultured alone or co-cultured with a bacterium. *Penicillium* and *Scopulariopsis* species contributed the largest number of significant features, which is corroborated by their biosynthetic potential as these genera were found to have a high number of predicted biosynthetic gene clusters via antiSMASH (BGCs; **Figure 1**; **Figure S2**)^27^. Conversely, *Fusarium*, the yeasts, and bacteria do not appear to contribute metabolites that could be detected via LC-MS and this likely is the reason for the lack of discrete clustering of those samples. The fungal metabolomes did not appear to change in any significant manner when they were co-cultured with either of the two bacterial partners. This strong metabolomic influence of fungi relative to bacteria in the cheese rind system is in agreement with several previous studies where fungi had stronger impacts on the system compared to bacteria^20,28–35^. Pierce *et al*. found that bacterial fitness was impacted by fungal growing partners whereas statistical analysis of *P. psychrophila* JB418 and *E. coli* K12 metabolomic profiles were not fruitful because bacterial features were simply not present in co-culture feature lists^21^.

### Distribution of Annotated Metabolites in Bacterial-Fungal Co-Cultures

In addition to the global analysis of metabolomic profiles, the feature list was processed to identify any known metabolites, analogues of known metabolites, and *in silico* predicted metabolites such as peptides and how they were distributed across samples. Biological replicates were pooled to represent a sample average and the resulting metabolomic profiles were organized into molecular networks generated using GNPS classical molecular networking (**Figure S3A**)^36^. Very few (7% of all nodes) were found to be unique to co-culture conditions and could be assigned as bacterial, fungal, or microbial as they were found only when species were grown together.

In the overall molecular networks of all extracts (bacterial, fungal, and co-culture), most metabolites were produced by fungi and were found in interactions with both *E. coli* K12 and *P. psychrophila* JB418 or ambiguously assigned as a general microbial metabolite. A large number of peptides were not species-specific as they were found in all cultures containing the filamentous fungi studies. This likely reflects the breakdown of casein or other cheese proteins by fungal proteases rather than production of specialized metabolites. The distribution of the classes of fungal metabolites reflected the patterns of both general and specific fungal effects seen previously in the RB-TnSeq data^21^. Since fungi produce a greater number of metabolites (**Figure S3A**), they are likely to have a greater total effect on interactions with bacteria^37^. Fungal metabolite abundances provide an important consideration because microbiome studies are often focused on bacteria. If fungi are the primary drivers of microbial interactions, many of these interactions would be missed through omission of fungi from these studies. However, it is unclear as to whether the difference in specialized metabolite production is a product of the different biosynthetic potentials between fungi and bacteria or simply due to the fact that fungi have been found to be more abundant in cheese rind microbiomes with the exception of washed cheese rinds^18^. Therefore, it is important to understand whether the metabolite distribution observed was a function of the community composition and chemical extraction methods used or whether fungi dominate regardless of these factors.

### Bacterial Community Composition Enacts Negligible Changes to Penicillium Metabolomic Profiles

One reason for the limited effect of the bacteria on the metabolomic profiles above may be that the two bacteria make a limited suite of exometabolites (**Table S1**). To explore this, we conducted experiments with two Actinobacteria from the cheese rind system since Actinobacteria are generally considered high producers of specialized metabolites^20,38^. We evaluated the effects of fungal co-culture with Actinobacteria and compared the metabolite profiles to fungal co-cultures with Gammaproteobacteria by performing pairwise co-cultures of *P. solitum* #12 with either (1) *P. psychrophila* JB418, (2) *E. coli* K12, (3) *G. arilaitensis* JB182, or (4) *B. linens* JB5 in addition to their respective monocultures. Here, *P. solitum* #12 was chosen as a representative filamentous fungi as it has been shown to possess high biosynthetic potential and is a prominent producer of specialized metabolites (**Figure 1; Figure S2**)^21^. *G. arilaitensis* JB182 and *Brevibacterium linens* JB5 were chosen using IDBac for strain prioritization (**Supplemental Information, Dataset 3**)^24^. Sparse partial least squares discriminant analysis (sPLS-DA) revealed that bacterial monocultures tended to cluster closer to the media controls (10% CCA). On the other hand, monocultures and pairwise co-cultures with *Penicillium* appeared to cluster distinctly from the controls and bacterial monocultures; finer differences in *Penicillium* clustering such as differential abundance of various features appeared to be the result of differences in bacterial growth partners (**Figure 2**). This finding was recapitulated upon performing hierarchical clustering of metabolomics profiles, where metabolomic profiles from bacterial monocultures cluster with the media and metabolomic profiles derived from *Penicillium* monoculture and co-cultures clustered together (**Figure 3**). Co-cultures that contained *Penicillium* and *Pseudomonas* produce metabolomic profiles that were distinct from other *Penicillium* associated profiles. Further, the *Pencillium-Pseudomonas* metabolomic profile clustered between fungal profiles and the media profiles, indicating there may have been an unusually large amount of media related features in those extracts relative to other profiles. Overall, despite the change in community composition to include more “talented” producers of specialized metabolites, metabolomic profiles were still found to be primarily composed of fungal features. Next, different culture and extraction conditions were explored to determine their effects on detected community metabolite distribution.

**Figure 2:**
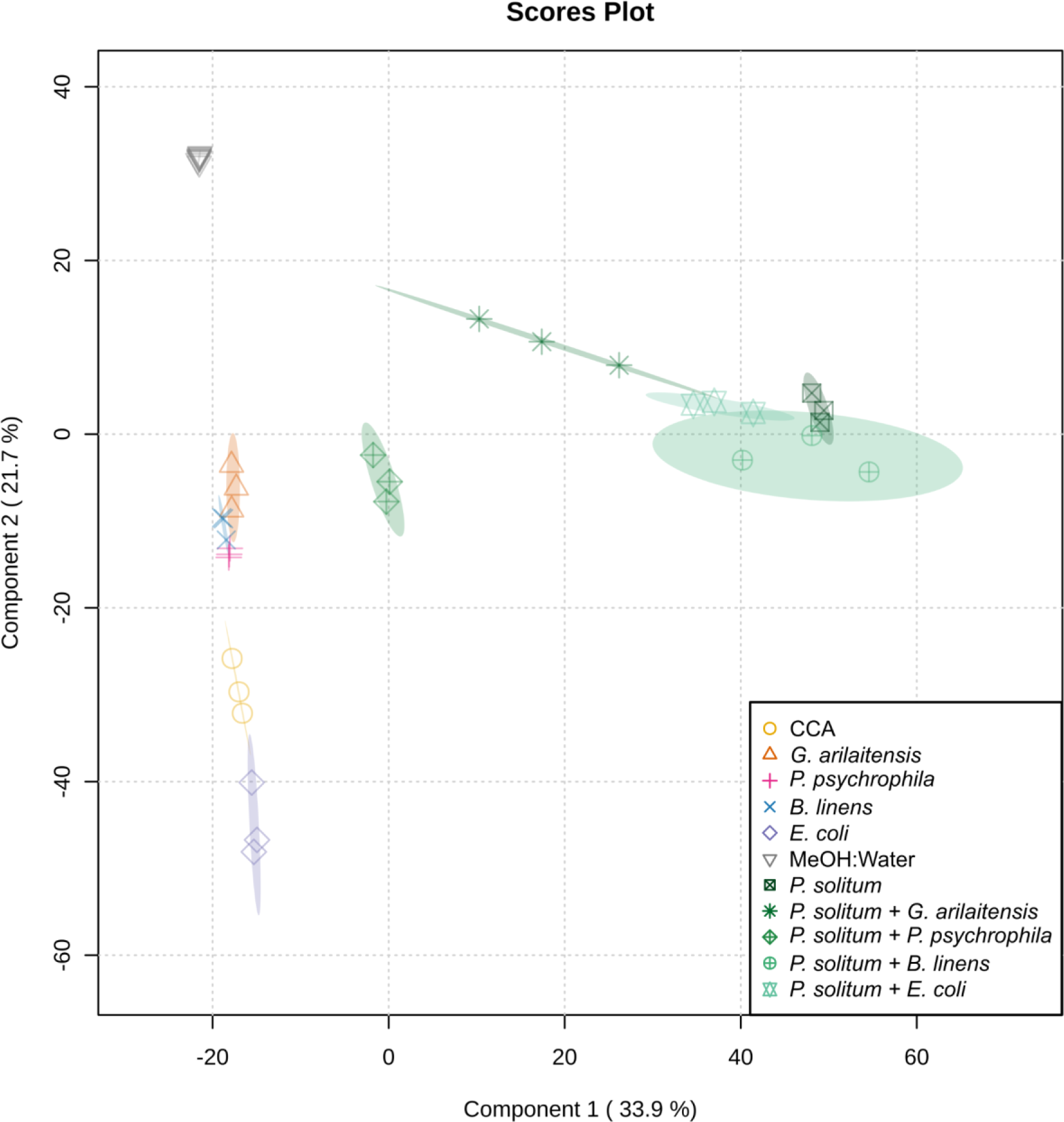
sPLS-DA plot showing solvent and media blanks, bacterial and *P. solitum* #12 monocultures, and bacterial-fungal pairwise co-cultures grown in the dark. sPLS-DA was performed with five components and five-fold cross-validation using the top 1000 features.

**Figure 3:**
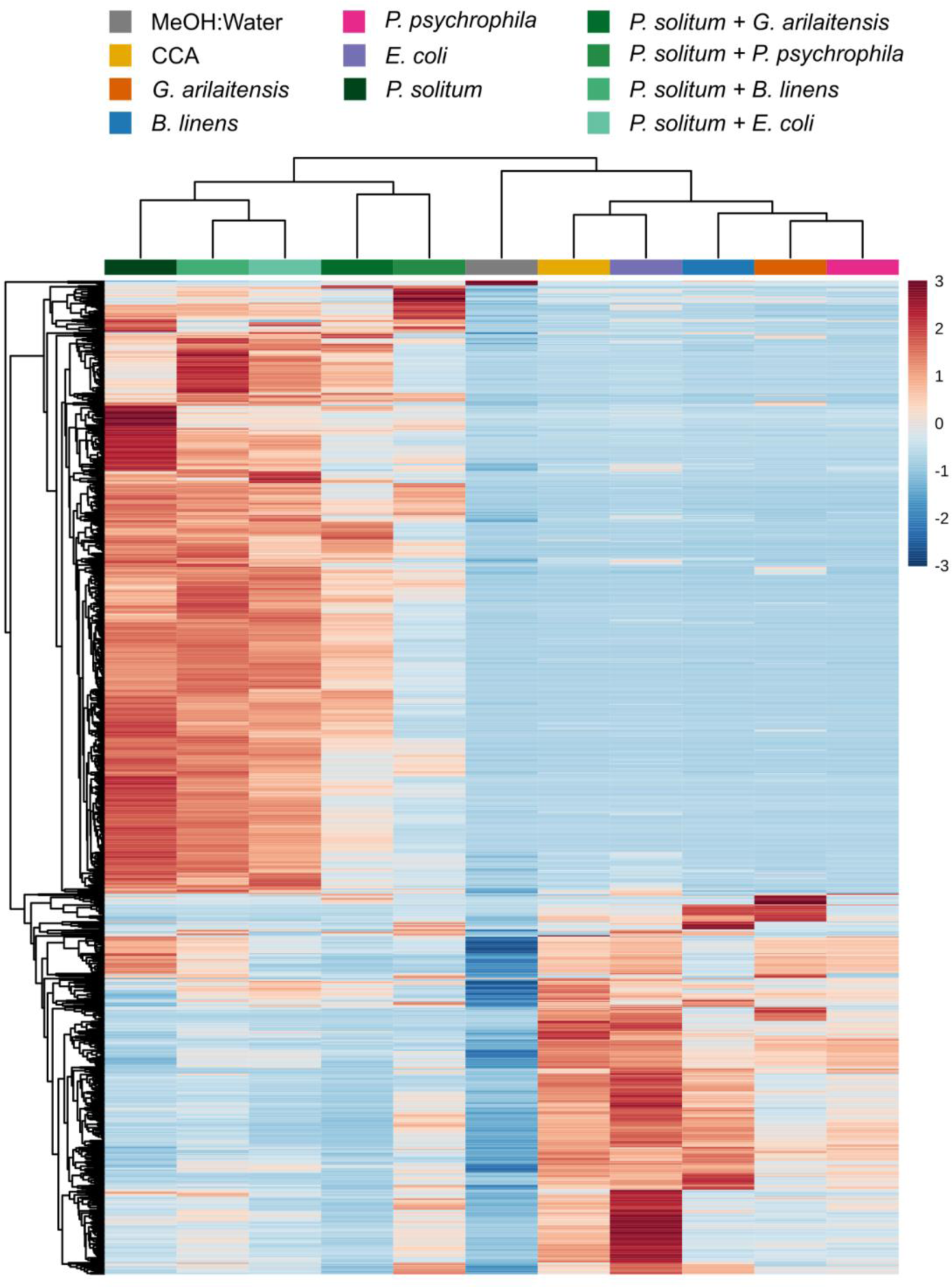
Hierarchical clustering of features and samples is displayed in this heatmap. Cultures grown here were grown in the dark. The top 1000 most significant features as determined by ANOVA are plotted by intensity in the sample, and only group averages are shown here with a group referring to the average of all biological replicates across one sample condition. Sample and feature clustering was performed using the unweighted pair group method with arithmetic mean (UPGMA) algorithm based on Euclidean distance.

### Sample Preparation is Increasingly Important When Complex Culture Media is Used

All conditions described above were grown on 100 mm diameter (∼31416 mm^2^) petri dishes, and during crude extract generation, all agar medium and microbial biomass were excised to perform solvent based extractions. However, typical fungal colonies can range from 30 - 50 mm in diameter (∼2827 - 7853 mm^2^), and typical bacterial colonies are ∼10 mm in diameter (∼314 mm^2^); a typical crude extract contains large amounts of media by area. While some media such as Luria-Bertani (LB) agar or International *Streptomyces* Project-2 (ISP-2)^39^ agar have inherently simple compositions, CCA is made from freeze dried cheese curd whose components carry over to crude extracts analyzed via MS as previously noted^22,40^. Therefore, rather than generating crude extracts from 100 mm samples, we next tested whether controlled, smaller areas excised from bacterial and fungal monocultures as 30 mm diameter plugs (i.e. the diameter of a 50 mL conical tube) could increase our ability to detect bacterial metabolites. These small plugs had a roughly ten-fold decrease in surface area (**Dataset 4**).

Based on the metabolomic profiles, despite the decrease in extracted surface area, bacteria continued to cluster with the media controls, while a majority of the features detected were of fungal origin (**Figure 4 & 5, Dataset 4**). The large cluster of upregulated features in the *Penicillium* monoculture appear to be fungal features that were most likely masked by the presence of media features in previous analyses.

**Figure 4:**
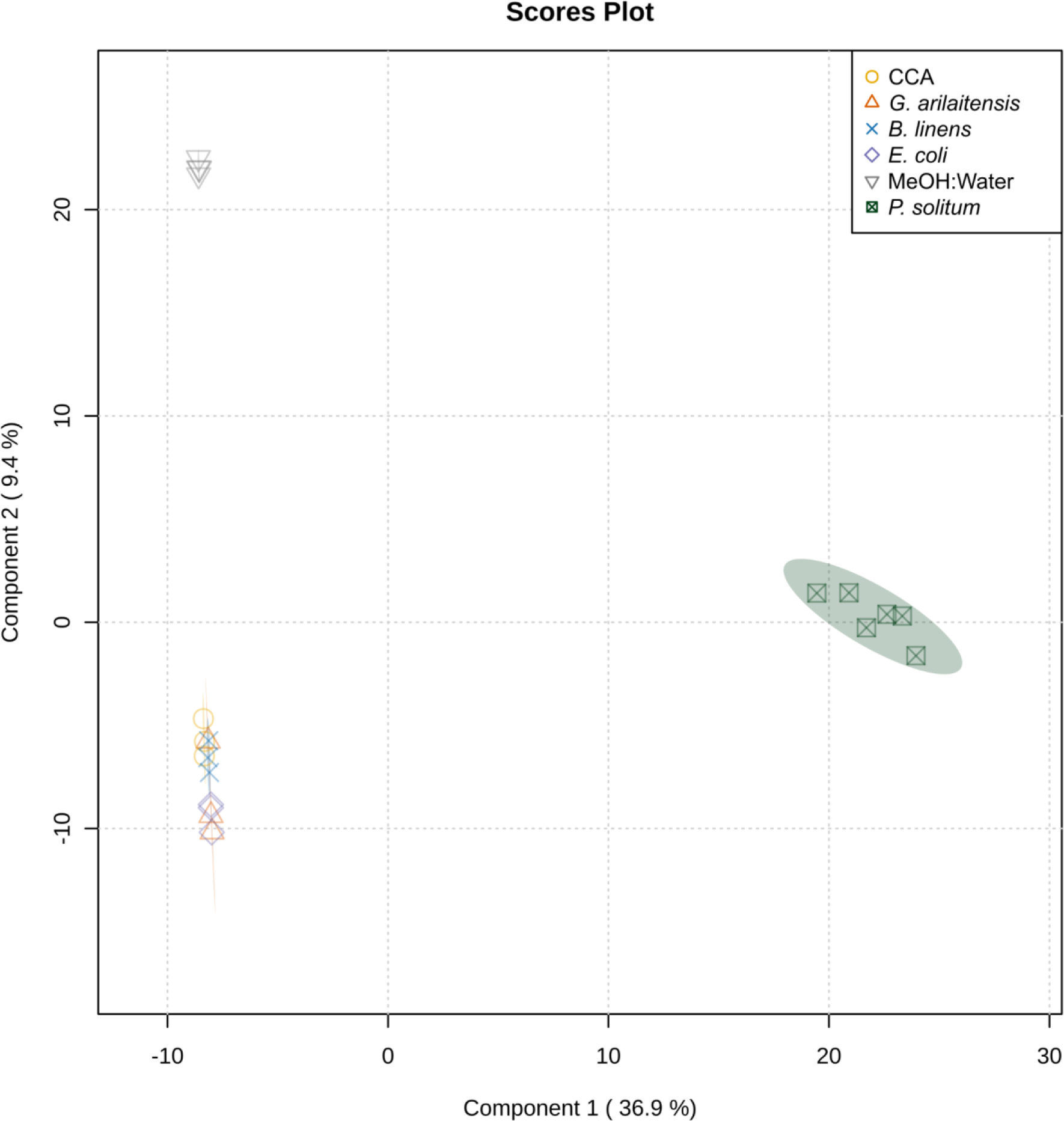
sPLS-DA plot showing solvent and media blanks, bacterial and *P. solitum* #12 monocultures grown in the dark in which chemical extracts were obtained from 30 mm plugs of each culture. sPLS-DA was performed with five components and five-fold cross-validation using the top 250 features. Here, a lower number of features was used since the inclusion of more features resulted in clustering based on run to run variation.

**Figure 5:**
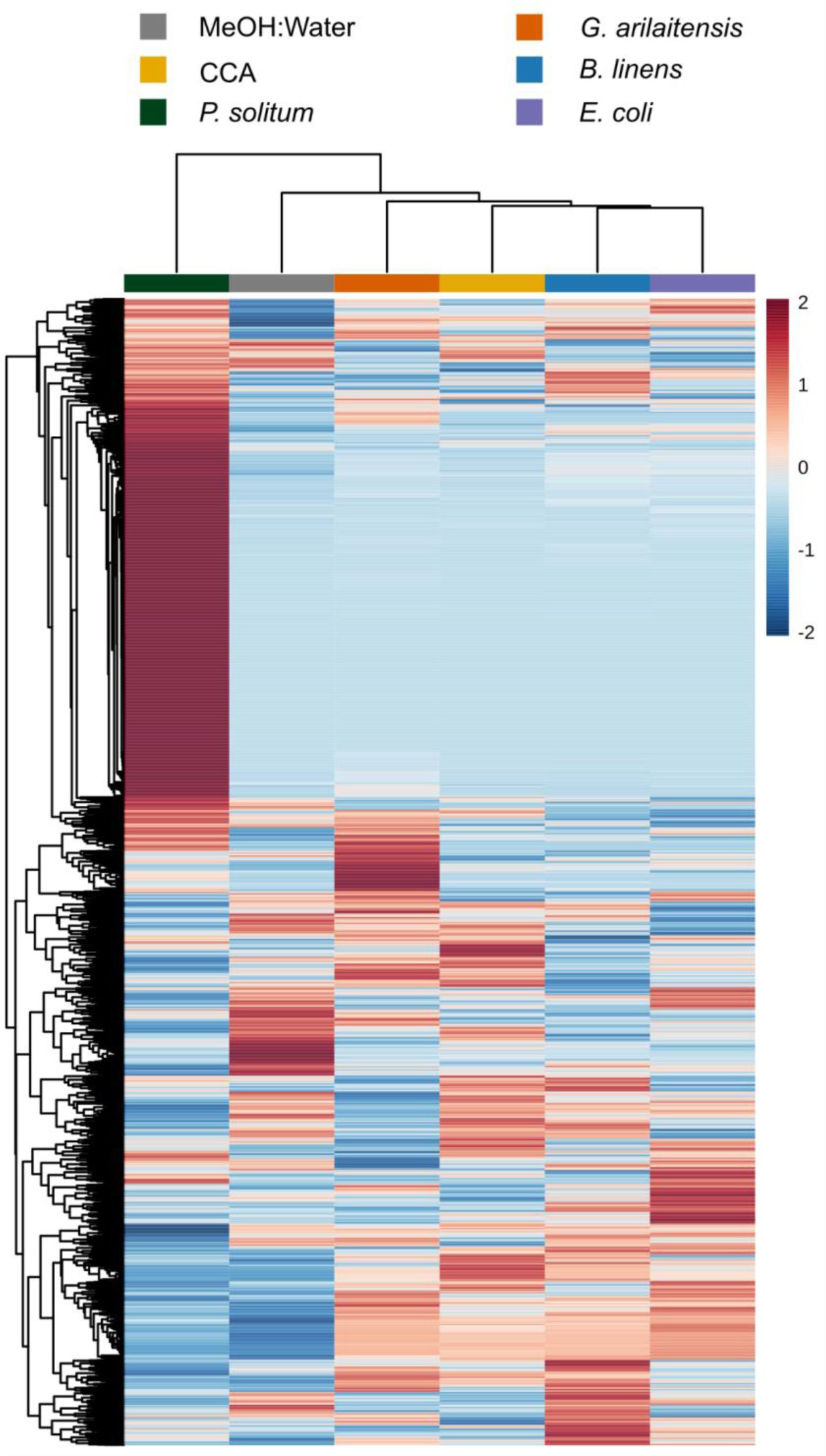
Hierarchical clustering of features and samples is displayed in this heatmap. Cultures grown here were grown in the dark. The top 1000 most significant features as determined by ANOVA are plotted by intensity in the sample, and only group averages are shown here with a group referring to the average of all biological replicates across one sample condition. Sample and feature clustering was performed using the unweighted pair group method with arithmetic mean (UPGMA) algorithm based on Euclidean distance.

GNPS feature based molecular networking, using the combined **Datasets 3** and **4** were processed via MZmine2, was used to annotate and analyze the samples^25,41^. Molecular networks are useful for visualizing the organization of MS/MS data into groupings based on structural relatedness^36,42^. These networks can identify analogues when unknown molecules group or cluster with known molecules from the GNPS database or commercially available standards. Additionally, GNPS also annotates features using library searches based on MS/MS similarity. Feature based molecular networking revealed the presence of fungal metabolites such as pyripyropene O, ML-236A, and cyclopeptine, with the latter two being new annotations present in Dataset 4 only (**Figure 6**; **Table S2**). While pyripyropene O was detected in both Dataset 3 and Dataset 4, there appeared to be a more than five-fold increase in abundance in Dataset 4 in the feature list generated through MZmine2. At the same time, ML-236A and cyclopeptine were only detected in the MZmine2 feature list from **Dataset 4**, indicating that this plug extractions were able to increase sensitivity for specialized metabolite detection (**Supplemental File**). Pyripyropene O was discovered from *Aspergillus fumigatus* FO-1289-2501 and previously shown to inhibit acyl-CoA cholesterol acyltransferase (ACAT) activity in rat liver microsomes^43^. ML-236A was first isolated from *Penicillium citrinum* and has been shown to be a hydroxymethylglutaryl-CoA reductase inhibitor that acts as an anti-cholesteremic agent^44^. Cyclopeptine is an intermediate in the biosynthesis of benzodiazepine alkaloids. Its presence here indicates the potential for *P. solitum* #12 to biosynthesize compounds of such a structural class such as sclerotigenin, which has previously been found in cheese derived *Penicillium* species, or the circumdatins^45–49^. Based on these results, crude extract generation from appropriately sized plugs improves feature detection for metabolomic analysis of microbial cultures.

**Figure 6:**
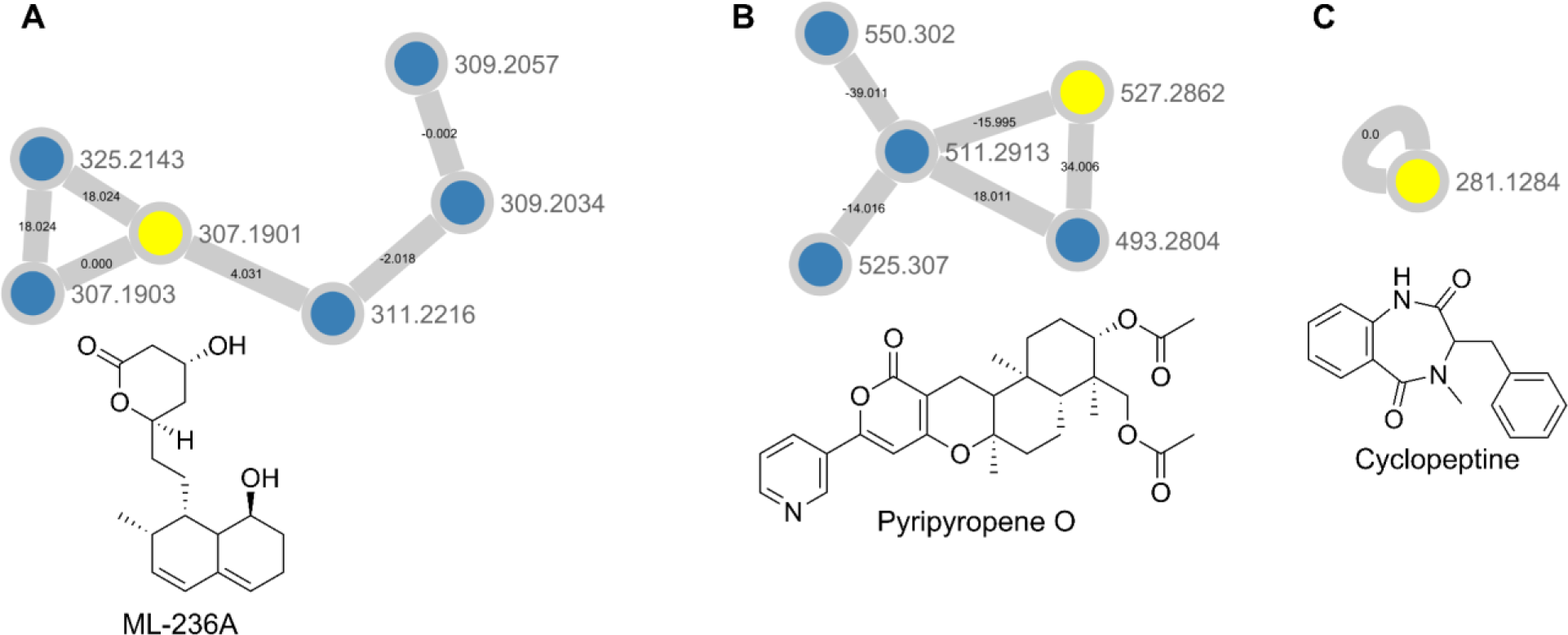
Molecular network showing (A) ML-236A, (B) pyripyropene O, and (C) cyclopeptine. Nodes highlighted in yellow indicate that it was a library hit in GNPS, while nodes in blue represent potential analogues detected in our dataset.

Lastly, we wanted to determine whether the removal of microbial biomass, and therefore removal of any colony-associated specialized metabolites prior to extraction, resulted in any changes to the resulting metabolomic profiles. Furthermore, we were interested in determining whether removal of the biomass resulted in any changes in metabolite distribution, as the fungal biomass often dwarfs the bacterial biomass (**Figure S4**). Previous work has shown that inclusion or removal of fungal biomass prior to performing imaging mass spectrometry caused changes in the features detected during acquisition; most features found to be colony associated could not be detected when the fungal biomass was removed, and most features found to be secreted could not be detected when the fungal biomass was present^50^.

*P. solitum* #12 was grown in co-culture with *E. coli* K12 along with their respective monocultures, and cultures were grown on PCAMS or on top of a nitrocellulose membrane placed on PCAMS (**Dataset 5**). Nitrocellulose membranes act as a barrier between the microbes and media while still allowing nutrients and specialized metabolites to diffuse freely, enabling the ability to completely remove the microbial biomass prior to crude extract generation while still maintaining secreted specialized metabolites. The use of membranes in the form of thin cellophane films has previously been demonstrated by Mascuch *et al*. and Larsen *et al*.^47,50^. However, as opposed to extracting the biomass above the membrane after culturing, only the media and secreted metabolites below the membrane were extracted. Clustering of the metabolomic profiles obtained from Dataset 5 showed that each culture condition tended to cluster with their counterparts grown on the nitrocellulose membrane, meaning the presence or absence of the microbial biomass made little to no difference in the makeup of the resulting metabolomic profile (**Figure S5 & S6**). This is most likely due to the fact that a majority of specialized metabolites observed in the cultures appear to be excreted; removal of the microbial biomass would only meaningfully change the profiles if intracellular specialized metabolites significantly contributed to the makeup of the crude extracts and resulting metabolomic profiles. Having optimized LC-MS/MS feature detection, molecular networking was used to annotate fungal metabolites and further explored for biological significance in the context of the cheese rind microbiome.

### Distribution of Annotated Fungal Metabolites

A molecular network of only fungal metabolomic features generated from Dataset 1 showed that the distribution of fungal metabolites reflected the pattern of both general and specific fungal effects that were previously observed at the genomic fitness level (**Figure S3B**)^21^. Through RB-TnSeq, species specific gene sets with enriched functions in pairwise co-cultures with yeast species were found to be smaller when paired with native cheese rind bacteria (**Table S3**); gene sets with enriched functions in pairwise co-cultures with yeast species were found to be similar to pairwise co-cultures with filamentous fungi (**Table S4**)^21^.

Overall, the *Penicillium* spp. and *Scopulariopsis* spp. account for the majority of identified metabolites (**Figure S7**). As well-known prolific producers of specialized metabolites, and with the high number of predicted biosynthetic gene clusters in the genome, *Penicillium* spp. collectively produce approximately 50% of the known metabolites in these co-cultures. While the bioactive metabolites identified in these analyses did not show significant changes in abundance upon co-culture, it is still likely that they are either directly or indirectly involved in these interactions.

Identified metabolites include those with known bioactivities in various systems, with the siderophore class of metabolites being found in five of the filamentous fungi (**Figure S7**; **Table S3**). Siderophores are small molecules that are produced to facilitate microbial uptake of insoluble iron. They accomplish this by forming complexes with Fe^+3^ with varying degrees of affinity for iron and other trace metals depending on the molecular structure and pH^51,52^. Previous work by Pierce *et al*. had shown that *E. coli* K12 fitness was dependent on expression of the *fep* operon containing genes that encode for native siderophore transport (i.e. enterobactin)^21^. In the absence of *fep* expression, the Fhu system was able to take advantage of hydroxamate-type siderophores produced by filamentous fungi^21^. All three of the *Penicillium* spp. had a library match to ferrichrome, and the *Penicillium* sp. and *Fusarium* networked with various forms of coprogen (i.e. coprogen, coprogen B, palmitoylcoprogen). Comparison of *Penicillium* and *Fusarium* extracts to ferrichrome and coprogen B standards provided Level 1 identification of these compounds.

Manual inspection of the fragmentation validated the clustering of coprogen analogues with varying carbon chain lengths and double bonds with low ppm error to provide level 2 identification^21^. Based on these data, the importance of siderophores in BFIs warranted further experimentation to explore and elucidate siderophores produced by cheese rind derived filamentous fungi.

### Siderophore Detection in Filamentous Fungi

The chrome azurol S (CAS) assay was performed on fungal extracts to unambiguously detect the presence of non-chelated siderophores that were not immediately apparent in the MS analyses (**Figure S8**). Importantly, the CAS assay is only capable of indicating the presence of non-chelated siderophores. Therefore, siderophores that are already complexed with iron will not produce the colorimetric change associated with iron binding. All of the filamentous fungi were positive by CAS assay for the presence of siderophores while the yeasts were negative. Therefore any further siderophore analysis was only performed on the filamentous fungal species.

Previous studies have identified unknown siderophores by employing liquid chromatography-inductively coupled plasma-mass spectrometry (LC-ICP-MS), which combines chemical separation of compounds with elemental analysis rather than analysis of intact protonated molecules which can disrupt iron chelation through the use of acidic modifiers common in LC-MS/MS analyses^53^. We utilized this hyphenated technique for siderophore analysis, which allowed for the direct measurement of elemental iron and other trace metals using LC-ICP-MS rather than relying on detection via ionization of the siderophores themselves (Dataset 6).

In order to observe iron chelation at a physiologically relevant pH, mobile phase solvents were adjusted to a pH of 6.5 prior to LC-ICP-MS analyses which ensured favorable conditions for iron chelation. At low pH, catecholates and hydroxamates (common functional group motifs on siderophores) are fully protonated and less likely to chelate iron^52^. Biological replicates of extracts were pooled by combining equal amounts of each replicate to create one pooled sample representing the average of all replicates. Solvent and media extract controls were analyzed to establish baseline signals for trace metals. The CCA media contains non-specific iron binding complexes such as phosphates, oxalic acid, citric acid, and various proteins^54–56^. Iron bound to these media-derived biomolecules produced two prominent peaks in the chromatogram (**Figure 7**). An iron(III) chloride solution (FeCl_3_, 10 µM) was used to determine where unbound iron eluted, and the CCA was treated with the FeCl_3_ to identify non-specific iron chelation not related to the presence of microbial siderophores.

**Figure 7:**
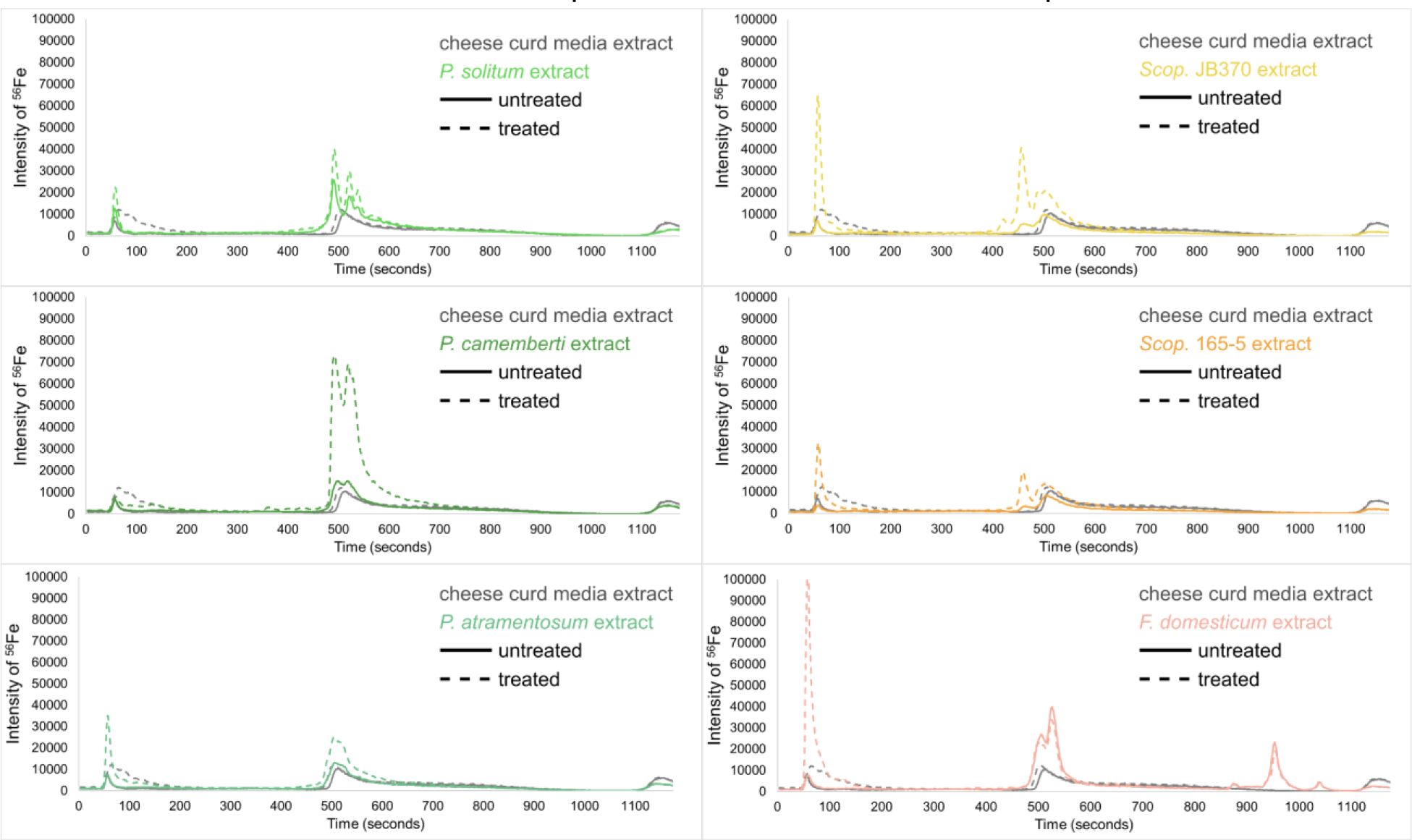
LC-ICP-MS chromatograms of fungal monocultures with FeCl_3_. Traces are colored according to fungal species and dotted lines represent treated extract traces. An increase in peak from treated to untreated indicates the presence of non-chelated siderophores in the extracts.

In fungal monocultures, a large increase in the intensity of ^56^Fe with different peak profiles and retention times indicated the presence of specific iron-binding complexes (siderophores; **Figure 7**). Only the filamentous fungi (*Penicillium, Scopulariopsis, Fusarium*) are shown because the yeast species did not show any differences in their LC-ICP-MS peak profiles compared to the controls coupled to the initial observation that they did not appear to contain any non-chelated siderophores in the CAS assay. Based on these LC-ICP-MS chromatograms, all of the filamentous fungi have iron-bound siderophores present in their respective extracts. To test whether some of these or other metallophores may be complexed with other trace metals, isotopes of other transition metals were also monitored including ^48^Ti, ^55^Mn, ^57^Fe, ^59^Co, ^60^Ni, ^63^Cu, ^64^Zn, ^65^Cu, ^66^Zn, and ^95^Mo. There was no evidence in the other channels for binding any of these metals and only peaks for ^56^Fe and ^57^Fe were observed in all the analyses performed on these extracts. As the most highly abundant naturally occurring isotope, ^56^Fe produced the most intense peaks and thus only data for ^56^Fe is shown. These results were inline for all fungi and detected siderophores with the exception of *Scopulariopsis* sp. JB370. This strain did not have any detected siderophores in the GNPS networks, however AntiSMASH analysis of the *Scopulariopsis* sp. JB370 genome predicted a biosynthetic gene cluster for dimethylcoprogen, but manual searches in the raw data for this metabolite and similar analogues did not yield any matches, indicating the potential presence of a cryptic BGC.

### Identification of A Novel Coprogen Analog from Scopulariopsis sp. JB370

Based on the LC-ICP-MS evidence of siderophores in the *Scopulariopsis* extracts, the LC-ICP-MS unique peak profiles and LC-MS/MS was reconducted under matching conditions to the LC-ICP-MS experiments (Dataset 7). *Scopulariopsis* extracts were treated with FeCl_3_ to saturate siderophores for detection of the distinct isotopologue pattern produced by the presence of iron. A comparison of treated and untreated extracts showed an increase of *m/z* 836 upon treatment with iron (**Figure 8A**). Inspection of peak profiles showed a concurrent decrease in *m/z* 783 which would represent the unchelated form (**Figure 8B**), and retention time shifts reflected the phenomenon of metal chelation shifting retention times to slightly more polar elution^57^. Additionally, *m/z* 836 exhibited a small peak 2 Da preceding the monoisotopic peak, representing the natural abundance of ^54^Fe (**Figure 8C**). Fragmentation patterns suggest a coprogen analogue with the presence of a terminal carboxylic acid (**Figure 8D & 8E**). A carboxylic acid motif could be found on previously discovered hydroxamate type siderophores such as basidiochrome and ferrichrome A^58,59^. However, the carboxylic acid would represent a new motif in the coprogen class of siderophores. Subsequent attempts to isolate quantities of this analogue for NMR analyses proved unsuccessful due to the apparent lack of further production by *Scopulariopsis* sp. JB370; EICs for *m/z* 783.3744 and *m/z* 836.2867 for these new extracts showed that both the free and iron bound forms of the novel siderophore were not present. Considering that fungal secondary metabolism has been reported as highly dependent upon growth factors such as sugar concentration in the medium, the issue may lie in the inherent variability of a complex media derived from an undefined matrix, in this case cheese curds^61,62^. This limited our level of identification to Level 2b, which is defined by Schymanski *et al*. as a “case where no other structure fits the experimental information, but no standard or literature information is available for confirmation”^63^. A synthetic route would need to be devised to fully confirm this structure.

**Figure 8:**
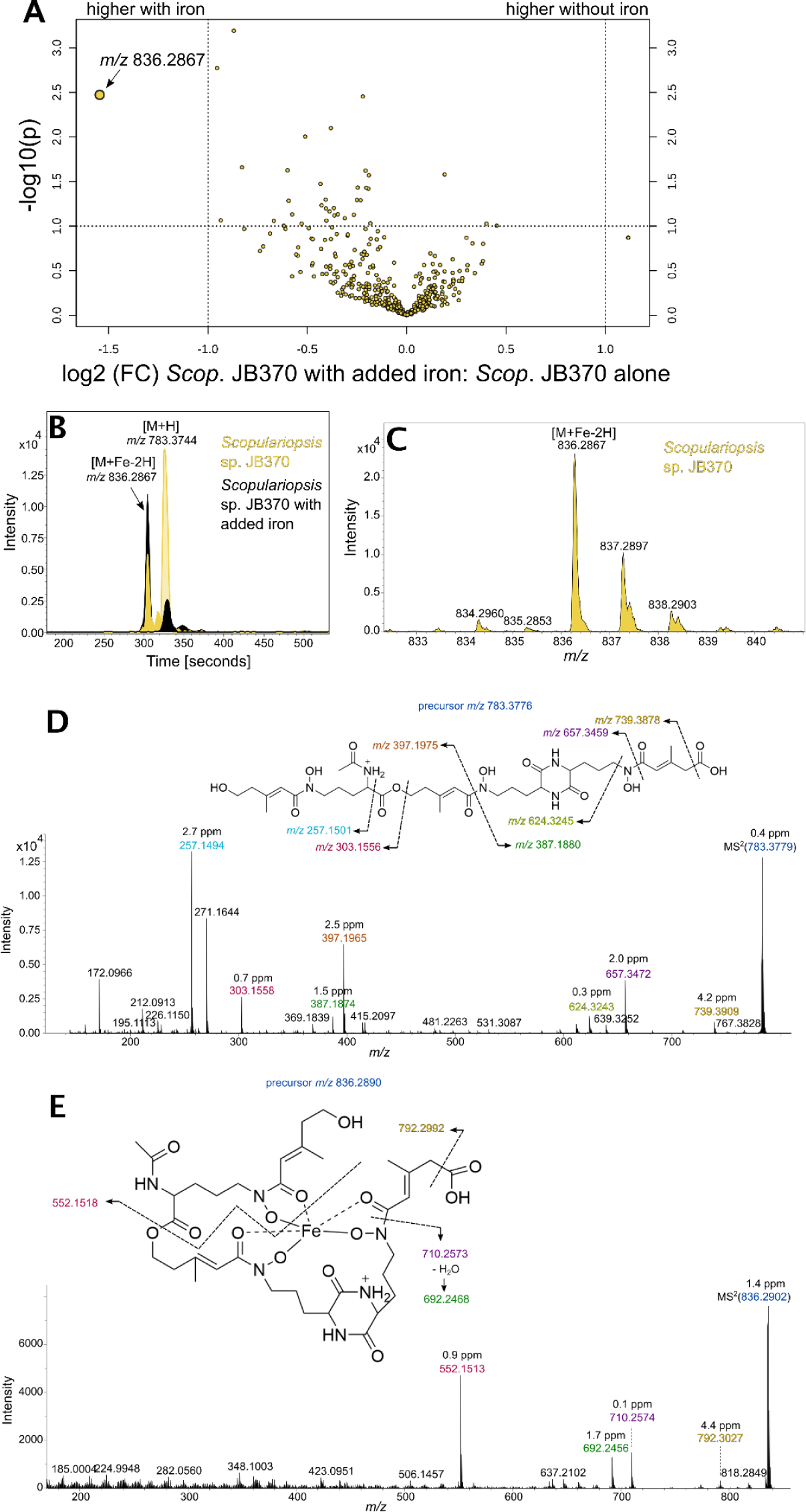
(A) Volcano plot comparing treated and untreated extracts from *Scopulariopsis* sp. JB370. *m/z* 836.2867 is upregulated in extracts treated with iron. (B) Comparison of EICs for *m/z* 783.3744 and *m/z* 836.2867 in treated and untreated extracts displays an inverse relationship in their abundance dependent on treatment with iron. (C) *m/z* 836.2867 displays an isotopic pattern characteristic in siderophores that have sequestered iron. (D & E) Putative fragments for *m/z* 783.3776 and *m/z* 836.2890, respectively.

## Conclusion

A panel of bacterial-fungal pairwise co-cultures were profiled to detect and annotate specialized metabolites implicated in BFIs derived from the cheese rind microbiome. As previously noted, fungi are important members in natural and synthetic communities derived from the cheese rind microbiome, and the metabolomic data verifies the importance of fungi in complex cultures^19,28,29,35,64^. A variety of cultivation methods were tested which highlight the complexities of moving beyond monocultures, as Morin *et al*. have previously used RB-TnSeq to show that pairwise interactions are not representative of interactions found in more complex community co-cultures. The complexity associated with expanding synthetic communities increases exponentially, requiring the optimization of methods that are able to accurately profile them. We have come to the conclusion that limiting extraction size by area increases the ability to detect microbial based features from complex media and cultures. Lastly, a putative novel coprogen analog featuring a terminal carboxylic acid moiety was detected with a Level 2b identification. Previous studies have linked production of metal chelating metabolites to pathological processes, and iron limitation has been shown to lead to expression of virulence factors^19,65–70^. With the wide range of biological repercussions of siderophore production, and numerous studies that have described the ability of siderophores to coordinate the trace metals inside iron, it is difficult to generalize the biological roles of these molecules to different environments. Trace metals are essential, but toxic at high levels, and their concentrations strongly influence microbial growth^71^. Siderophore production and biological downstream effects may be dependent on the environment in which they are found, and thus their effects upon neighboring species could be highly dependent on the availability of trace metals in the environment^72,73^.

Our data is in line with reports that some fungal species are capable of production of more than one siderophore and begs the question of why fungi have evolved to have redundant functional molecules produced by more than one BGC^74^. It would therefore seem likely that slightly different roles are accomplished by different siderophores. In competition for nutrients, if other microbial species were proximal and were also scavenging iron, it may be beneficial for the fungi to produce a variety of siderophores such that not all solubilized iron forms can be used by environmental competitors^75,76^. Perhaps biosynthesis of one type of siderophore can compensate for use of another siderophore by outside species, or a suite of siderophores with a varying range of affinity for iron and other metals allows the fungi to finely tune acquisition of biologically important trace metals. The variability of roles that siderophores can take on in different environments makes these molecules central to model studies and highlight the importance of mimicking the natural environment as closely as possible.

Just as striking as this variety of specialized metabolites produced for a singular purpose was the sheer abundance of specialized metabolites produced by fungi when compared with bacteria. Directly comparing metabolomic profiles made it clear that fungi are responsible for the majority of secondary metabolites produced in this system, and it stands to reason that this is the case in most environments^77^. While bacteria tend to be the subject of focus in microbiome systems, perhaps rightfully so due to their prevalence as pathogens and direct ties to human health, the capacity held by filamentous fungi for production of biologically active metabolites may outweigh their supposed inability to effectively colonize the gut. The direct and indirect effects of fungal metabolites could mean that with brief or periodic exposure, fungi may exert more influence over their environment than previously recognized^78^. Microbiome studies that measure metabolites in addition to microbial abundances are likely to shed light on this distinct possibility.

## Acknowledgements and Funding Sources

This work was supported by the National Science Foundation (NSF) grant MCB-1817955 and 1817887 (LMS and RJD), the University of California, San Diego (UCSD) Center for Microbiome Innovation (ECP), the UCSD Ruth Stern Award (ECP), National Institutes of Health (NIH) Institutional Training grant T32 GM7240-40 (ECP), NIH Institutional Training grant T32-AT007533 (JCL, GTL), NIH Individual Fellowship F31-AT010418 (JCL), and NIH NIGMS R01 GM125943-02S2 (LMS), UCSC Startup funds, UCSC Dissertation year fellowship (GTL).

