## Supplemental Information for "Metabolomics of bacterial-fungal pairwise interactions reveal conserved molecular mechanisms"

**Table of Contents**

BLANKA2: an Updated Algorithm for Blank Subtraction in Mass Spectrometry of Complex biological Samples S3

Using IDBac and antiSMASH to Prioritize Alternative Bacterial Growth Partners S4

Equation S1. Formula used to score strains based on IDBac results S4

Figure S1. Graphic depiction of co-culture inoculation layout S5

Figure S2. Predicted biosynthetic gene clusters in select fungal species S6

Figure S3. Molecular networks generated through GNPS S7-8

Figure S4. Bacterial-fungal co-culture strains possess different amounts of biomass S9

Figure S5. sPLS-DA plot for nitrocellulose culture dataset S10

Figure S6. Hierarchical clustering heatmap for nitrocellulose culture dataset S11

Figure S7. Hierarchical clustering of the fungal metabolites putatively identified using GNPS S12

Figure S8. CAS assay with bacterial and fungal monoculture extracts S13

Figure S9. Pseudo-phylogenetic tree created in IDBac S14

Figure S10. Molecular association network (MAN) created in IDBac S15

Table S1. Scores for each group of strains as determined by IDBac S16

Table S2. Metabolites identified from fungal species S17-18

Table S3. *P. psychrophila* JB418 species specific gene sets with enriched function S19

Table S4. *E. coli* K12 species specific gene sets with enriched function S20

Table S5. Metabolites identified from *Actinobacteria* species S21-22

References S23

**BLANKA2: an Updated Algorithm for Blank Subtraction in Mass Spectrometry of Complex Biological Samples**

More recently, development of new tools for handling MS/MS data have led to the re-evaluation of the previously developed approach to blank subtraction. Here, BLANKA2 and its updated approach to performing blank detection and subtraction from input datasets with an output format that is supported by a variety of downstream analysis tools is described.

BLANKA2 is written in Python 3.8 as a command line tool for Linux that takes LC-MS/MS sample and blank datasets in mzML format. Importantly, BLANKA2 only acts on MS/MS spectra, as methods and processing pipelines highlighted above (i.e. MZmine3) are better suited to performing blank subtraction on $MS^1$ spectra. Blank detection is performed using cluster based methods and relies heavily on falcon, a tool that performs scalable clustering of large amounts of MS/MS spectra[1]. Clustering is performed in falcon by 1) binning and hashing spectral features, 2) constructing nearest neighbor indexes, 3) computing a pairwise distance matrix based on cosine distance, and 4) performing density-based clustering. Bittremieux *et al*. discuss the process in more detail and have also benchmarked falcon to compare its performance to similar clustering tools[1]. Following MS/MS clustering, MS/MS spectra from sample dataset files are removed if they cluster with MS/MS spectra that are found in blank dataset files. Since blank subtraction performance is directly tied to MS/MS clustering performance, users are able to pass parameters to falcon to ensure the correct settings for their dataset are used. Blank subtraction from sample files is performed using the *MzMLTransformer* provided by the psims API[2].

BLANKA2 is available at https://github.com/gtluu/blanka2 as a command line tool for Linux, where documentation on its installation and usage is also available. Due to falcon's use of the Faiss library, which lacks a Windows implementation, BLANKA2 is not available on Windows[3]. However, users may run BLANKA2 and other Linux based software in the Windows Subsystem for Linux environment.

**Using IDBac and antiSMASH to Prioritize Alternative Bacterial Growth Partners**

*Glutamicibacter arilaitensis* JB182 and various *Brevibacterium* and *Brachybacterium* sp. (total 16 strains) were selected for co-culture experiments. Niccum *et al*. have previously observed genotypic differences at the strain level in the cheese rind microbiome, which implies there are also differences in the biosynthetic potential of different *Brevibacterium* and *Brachybacterium* strains[4]. Furthermore, analysis of **Dataset 2**, containing LC-MS/MS data processed with BLANKA2, using Global Natural Products Social (GNPS) classical molecular networking showed that these species potentially made a variety of known specialized metabolites or structurally related analogues of known specialized metabolites (**Table S5**). The IDBac workflow was used alongside a modified rapid extraction sampling method to prioritize for cheese rind microbes producing unique chemistry[5]. Intact microprotein profiles and small molecule data collected via matrix-assisted laser desorption/ionization time-of-flight mass spectrometry (MALDI-TOF MS) was used to create a pseudo-phylogenetic tree and molecular association network (MAN), respectively (**Figure S9 & S10**). A two-step strain prioritization algorithm was then used to group strains based on their pseudo-phylogeny and score each strain based on the amount of unique and shared chemistry with other strains in the dataset (**Equation S1; Table S1**). *Brevibacterium linens* JB5 and *G. arilaitensis* JB182 were chosen due to having high scores in addition to slightly above average number of BGCs relative to the other Actinobacteria that were screened. All data here constitutes **Dataset 8**.

**Equation S1**: Formula used to score strains based on IDBac results.

strain score = ∑(weight of edges connected to strain node ** 2)

**Figure S1**: Graphic depiction of co-culture inoculation layout for alternating “t streaks” in pairwise microbial co-cultures. Here, fungal colonies are denoted by the green and white streaked colony, while bacterial colonies are denoted by the gray-beige streaked colony. Streaks were performed perpendicular to each other.

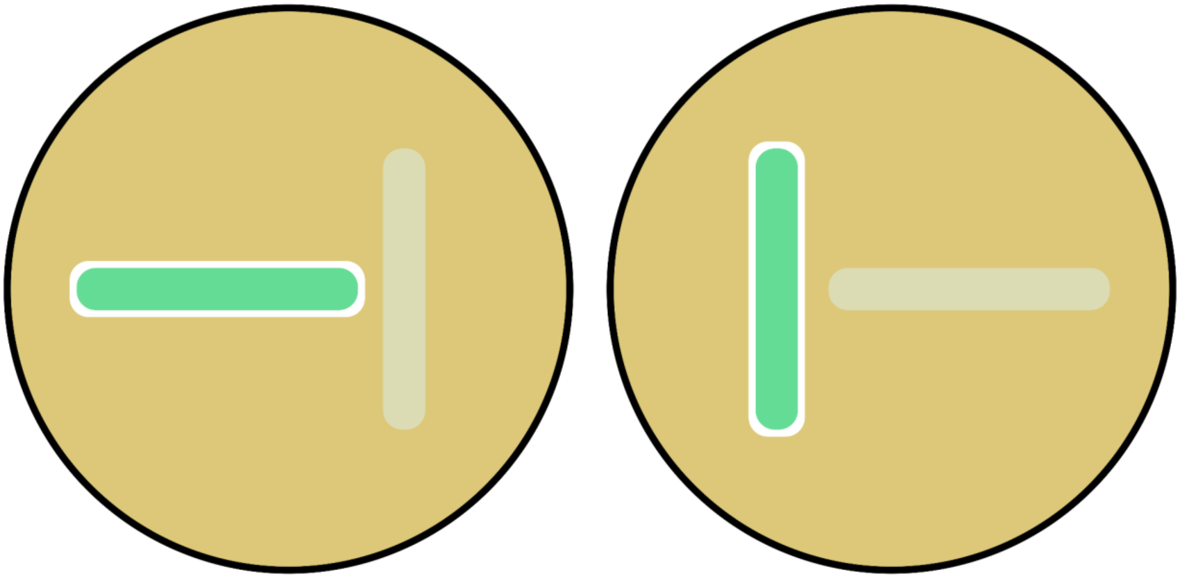

**Figure S2**: Predicted biosynthetic gene clusters (BGCs) found in select fungal species. Only fungal species included in this study with genome data available are shown here. The reported % known for each species refers to the % of BGCs represented here that have 100% similarity to BGCs that have been assigned to known metabolites. The *P. camemberti* genome is a publicly available complete genome and this likely accounts for the much higher number of identified BGCs. *P. atramentosum*, *P. solitum*, and *Scopulariopsis* sp. JB370 results are based on draft genomes compiled by the Dutton Lab.

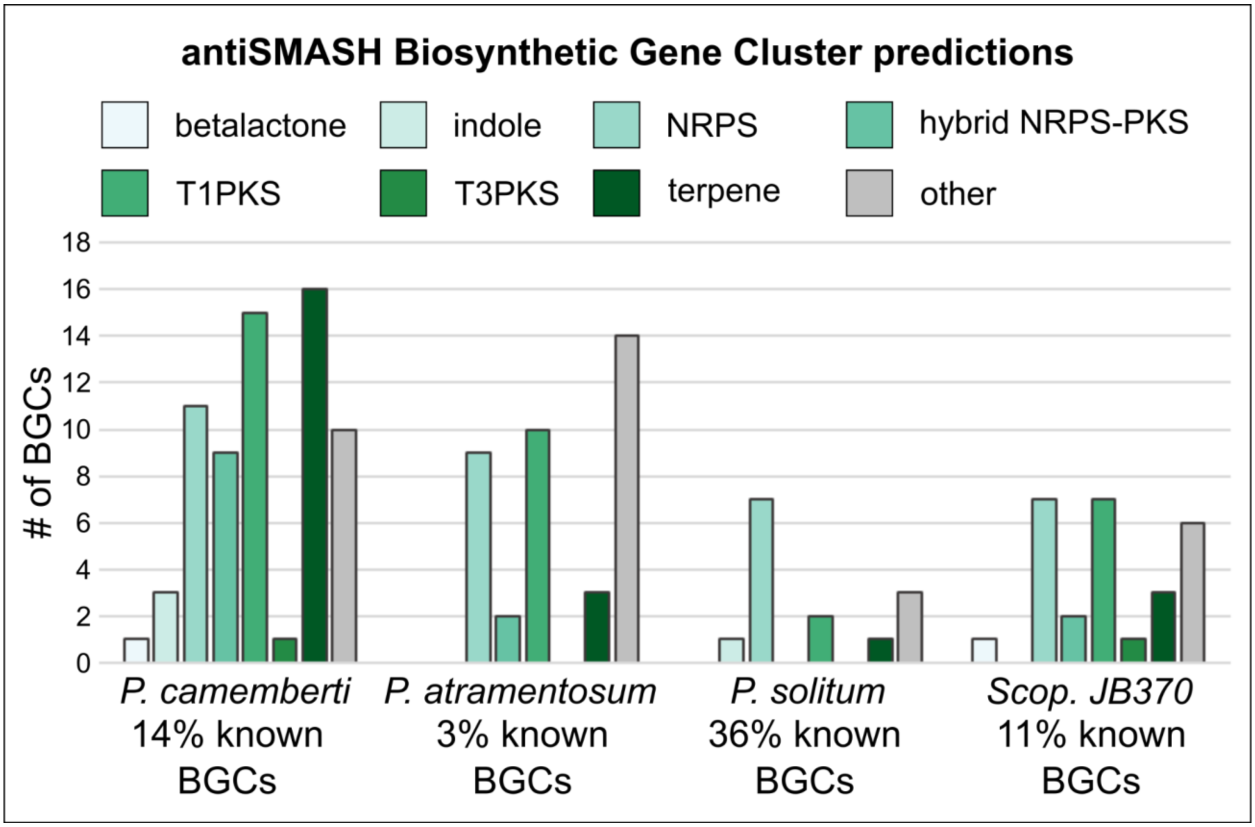

**Figure S3**: Molecular networks generated through GNPS of metabolites found in (A) all microbial cultures and (B) fungi derived metabolites. (A) The metabolites represented within the molecular network were assigned as either bacterial, fungal, or microbial according to the monocultures in which they were found. If they are present in fungal monocultures and co-cultures but not in bacterial monocultures, they are labeled as fungal and vice versa. If metabolites were found in both bacterial and fungal monocultures, they are labeled as microbial metabolites (represented as gray nodes). Network clusters containing different structural classes (i.e. amino acids, coprogens, etc.) can be identified. A majority of the nodes appear to be of fungal origin. (B) Features found in fungal monocultures are color coded to represent the specific fungal species in which they are present. The number of nodes found in the network that are specific to the yeast species suggests that their metabolites are not as diverse and abundant as those of filamentous fungi. Based on the structural classes of nodes found in different clusters, fungi are able to produce a variety of unique chemistry.

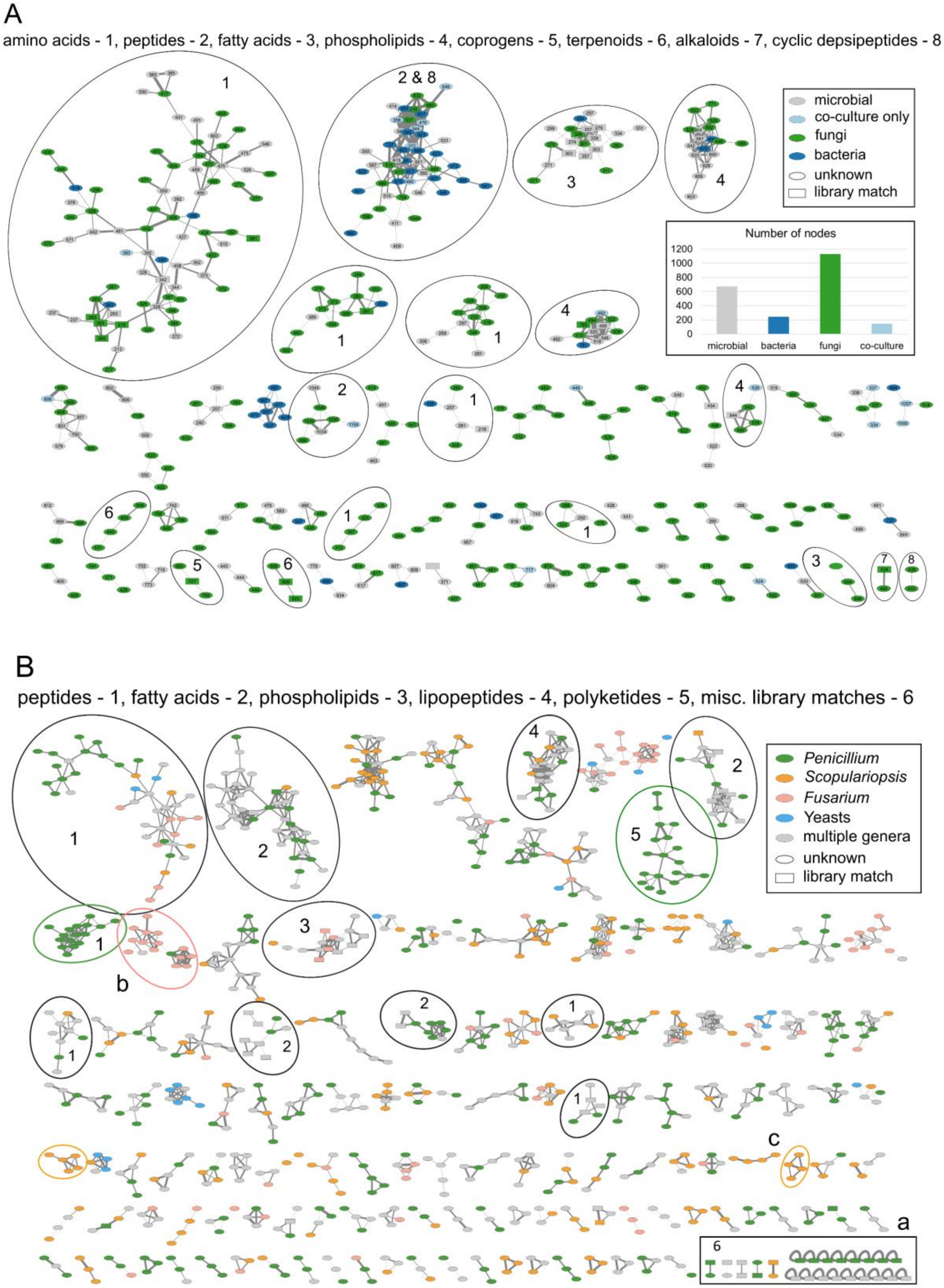
A
amino acids - 1, peptides - 2, fatty acids - 3, phospholipids - 4, coprogens - 5, terpenoids - 6, alkaloids - 7, cyclic depsipeptides - 8
microbial
co-culture only
fungi
bacteria
unknown
library match
Number of nodes
microbial
bacteria
fungi
co-culture
B
peptides - 1, fatty acids - 2, phospholipids - 3, lipopeptides - 4, polyketides - 5, misc. library matches - 6
Penicillium
Scopulariopsis
Fusarium
Yeasts
multiple genera
unknown
library match

**Figure S4**: Monoculture and pairwise bacterial-fungal co-culture images demonstrate the large inequality in colony biomass at 7 days of growth immediately prior to chemical extraction.

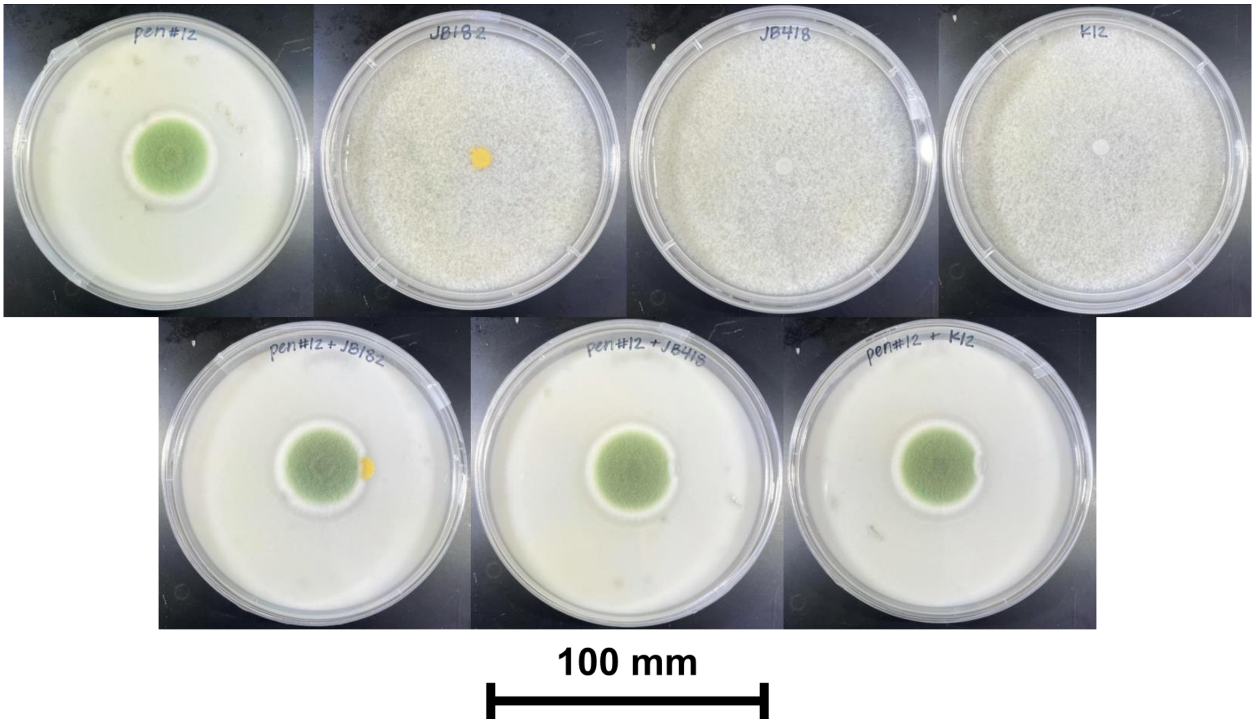

**Figure S5**: sPLS-DA plot showing solvent and media blanks, *E. coli* K12 and *P. solitum* #12 monocultures, and pairwise co-culture grown in the dark with and without the use of nitrocellulose filters. sPLS-DA was performed with five components and five-fold cross-validation using the top 1000 features.

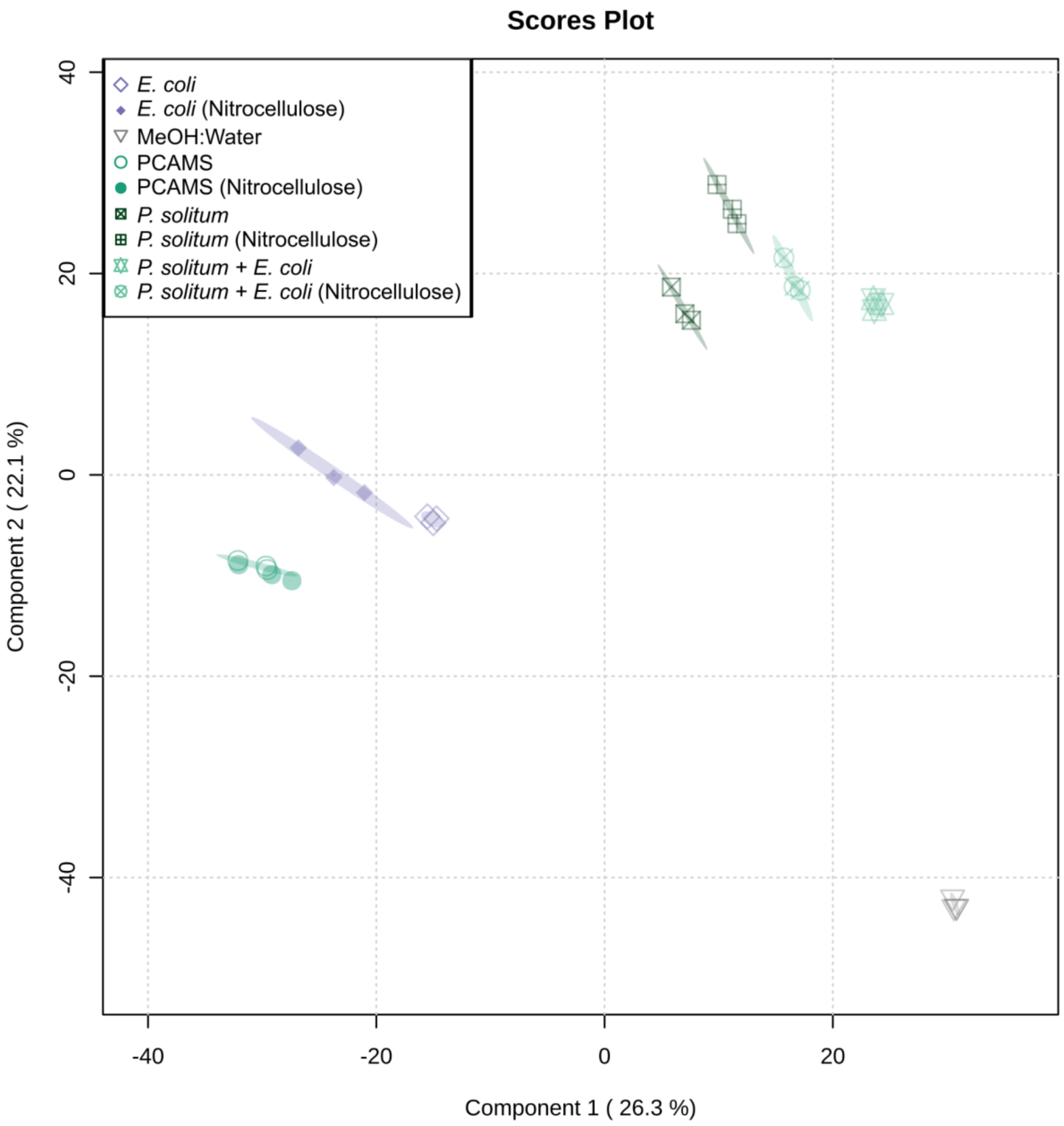

**Figure S6**: Hierarchical clustering of features and samples is displayed in this heatmap. Cultures grown here were grown in the dark. The top 1000 most significant features as determined by ANOVA are plotted by intensity in the sample, and only group averages are shown here with a group referring to the average of all biological replicates across one sample condition. Sample and feature clustering was performed using the unweighted pair group method with arithmetic mean (UPGMA) algorithm based on Euclidean distance.

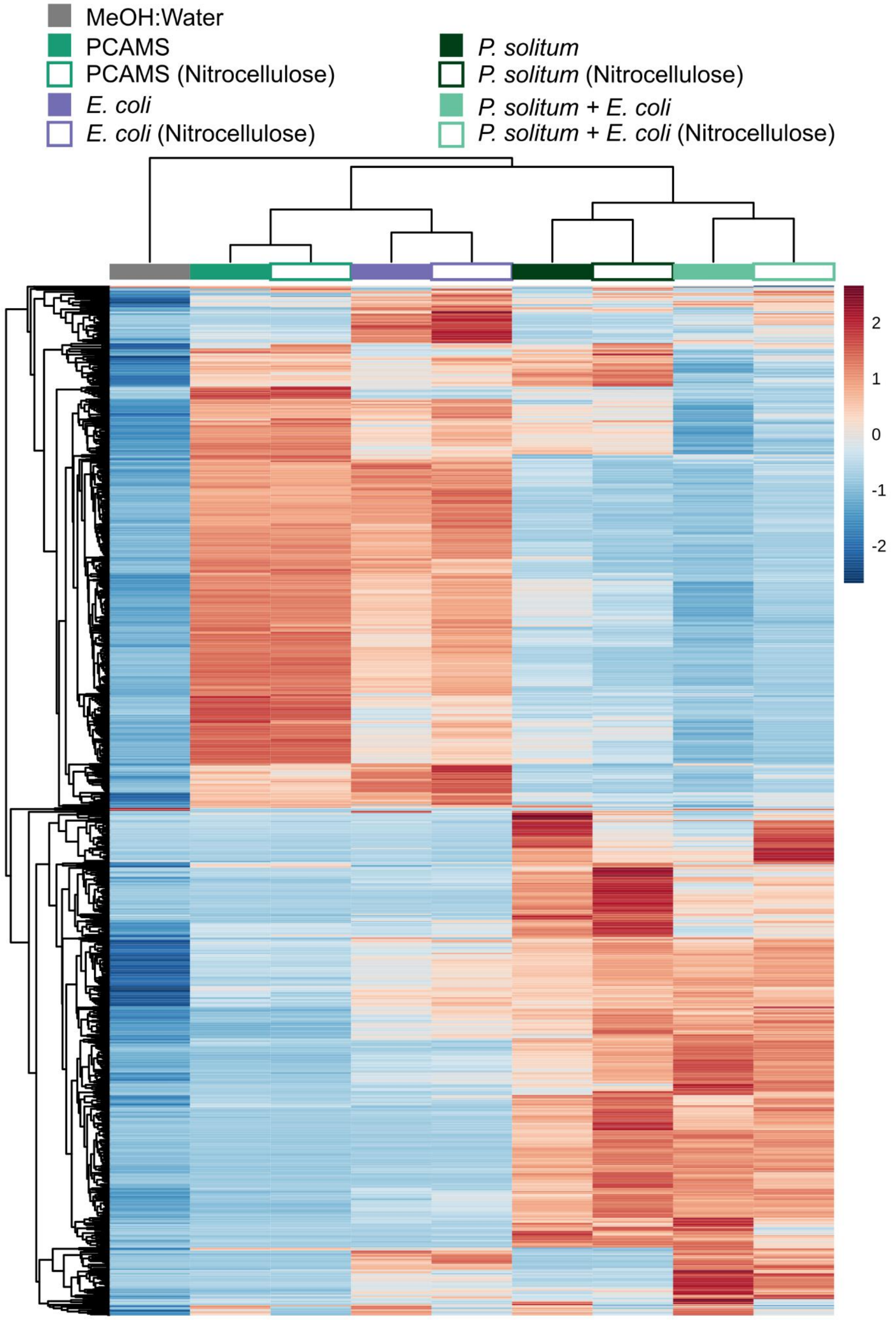

**Figure S7**: Hierarchical clustering of the fungal metabolites containing only features that were putatively identified using GNPS. *In silico* predictions are only included if fragmentation patterns from the raw data were indicative of the predicted compound class. For simplicity of visualization, features are shown here as a presence or absence map. The features have been parsed out to show different classes based on molecular structure (i.e. phenols, peptides) or known functions of molecules (i.e. siderophores). The pie chart lists approximate percentages of the total identified features described here as found in each culture of fungal monoculture.

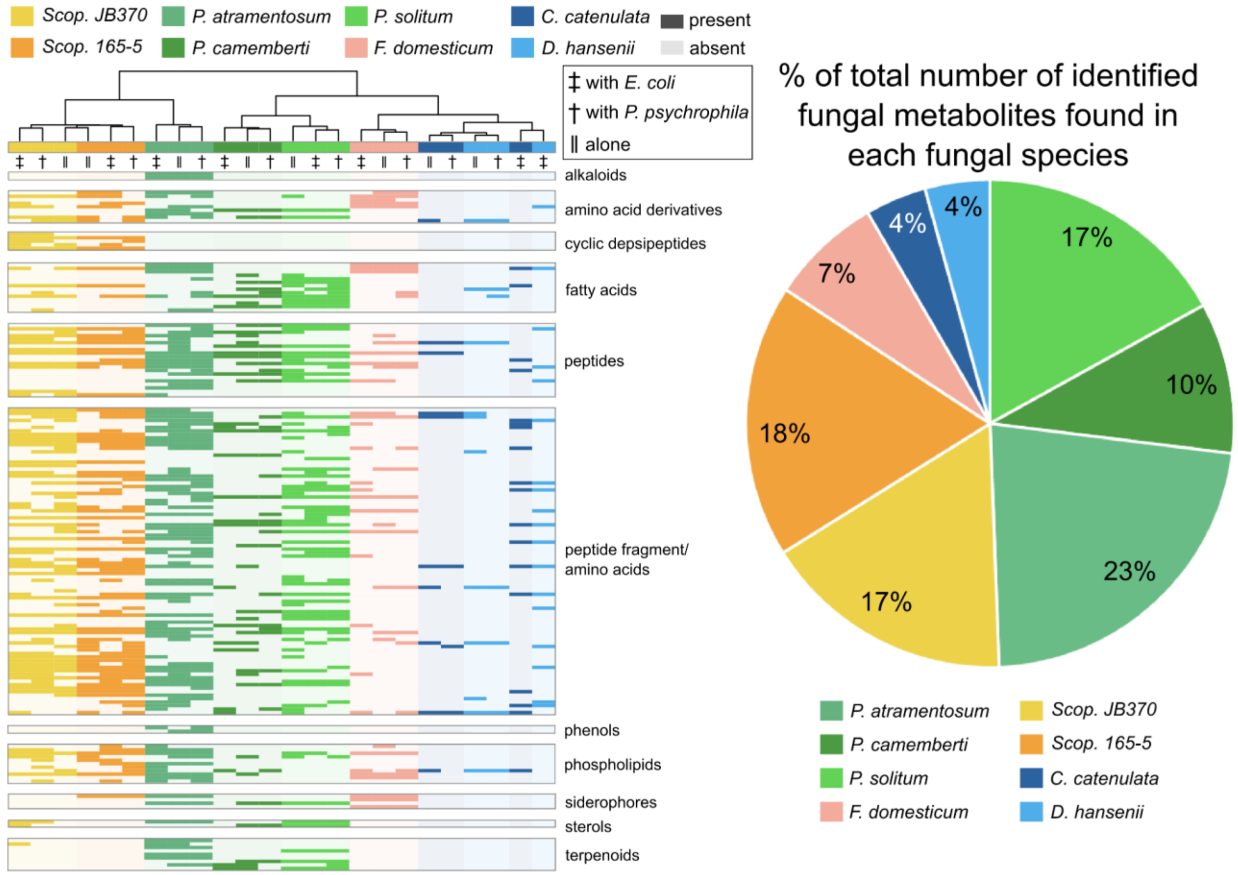

**Figure S8**: CAS assay with bacterial and fungal monoculture extracts. Deferoxamine standard was used for comparison in various concentrations and extracts were normalized to concentrations of 1 mg/mL and 0.1 mg/mL. * denotes a positive test result as indicated by visual inspection and $OD_{630}$.

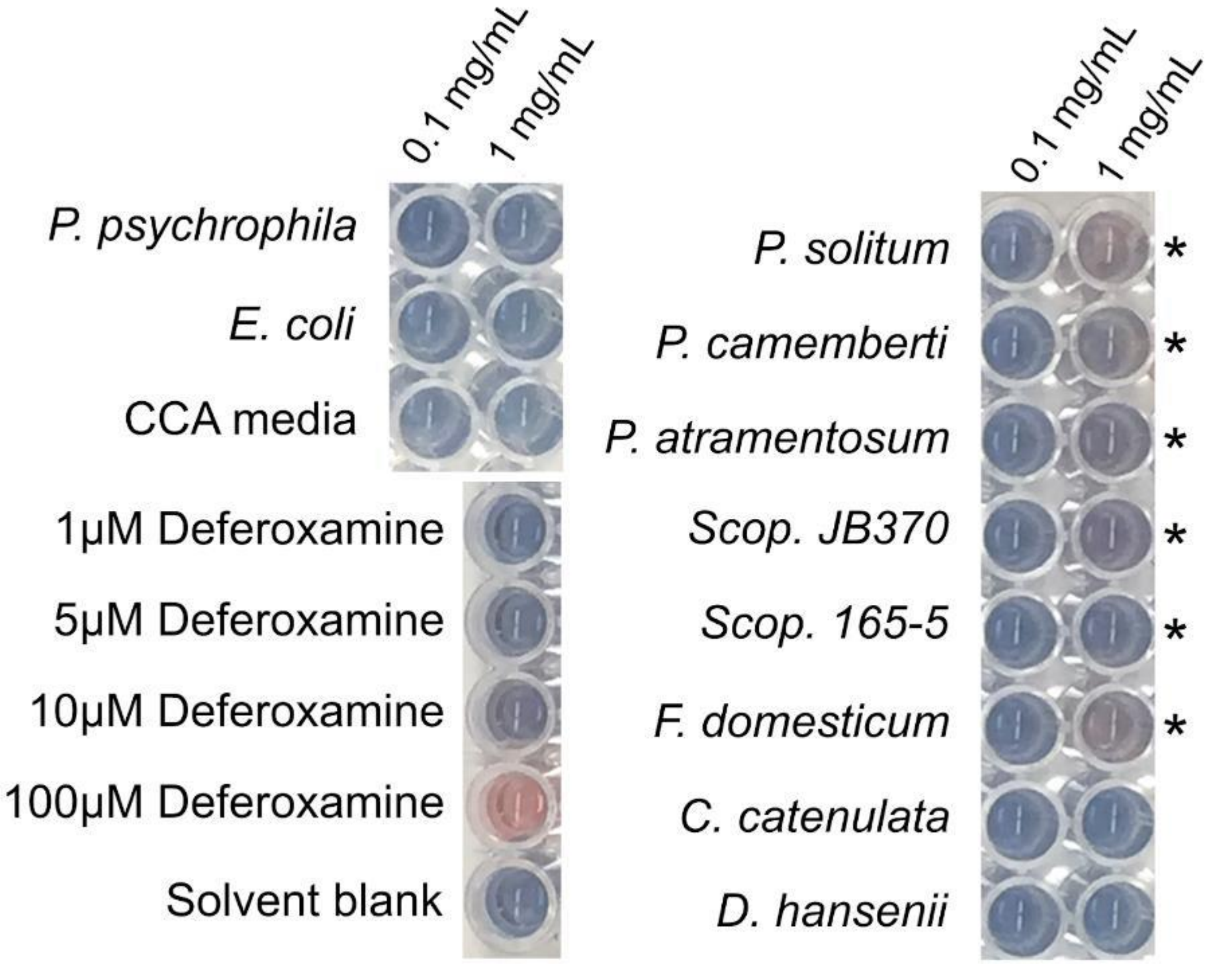

**Figure S9**: Pseudo-phylogenetic tree created in IDBac. Protein profiles for each strain were processed in IDBac and the dendrogram was created using peaks above a signal to noise ratio of 6 that were present in greater than 80% of replicate spectra (n = 4) in the mass range of *m/z* 3000 - 15,000. Hierarchical clustering was performed using cosine distances clustered by the unweighted pair group method with arithmetic mean (UPGMA) algorithm. At a tree height of 0.9, the dendrogram can be cleanly cut to yield two groups: one containing *Brevibacteria* and *Brachybacteria* and one containing all other bacteria.

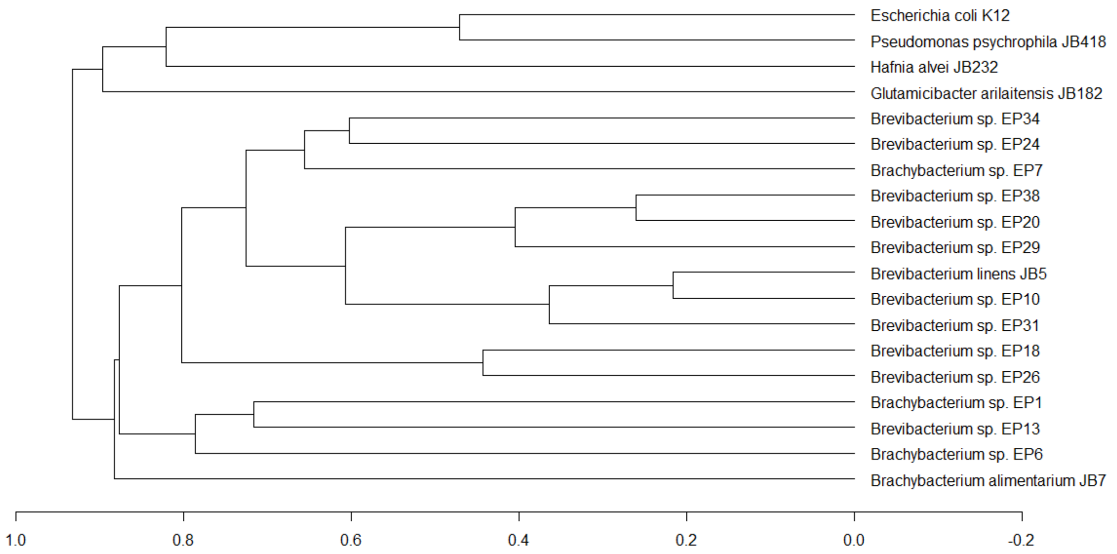

**Figure S10**: Molecular association network (MAN) created in IDBac. Small molecule profiles for each strain were processed in IDBac and the MAN was created using peaks above a signal to noise ratio of 3 that were present in greater than 80% of replicate spectra (n = 4) in the mass range of *m/z* 100 - 3000. Strains are color coded at the genus level. Here, *G. arilaitensis* JB182 and *E. coli* K12, being a couple of the most distant outgroups, are shown to produce the most unique chemistry in the dataset based on their vastly different biosynthetic capabilities relative to other strains analyzed here.

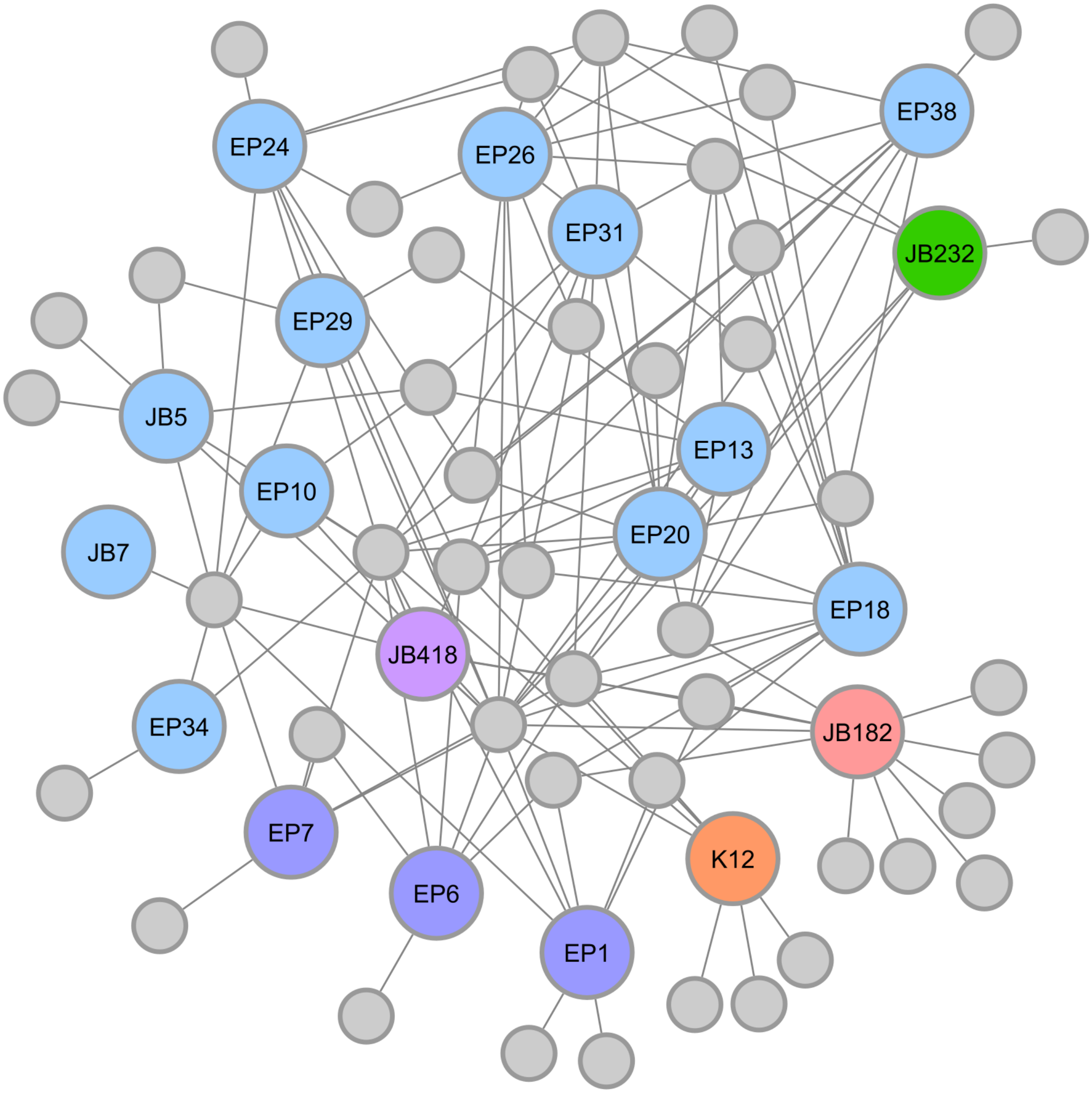

**Table S1**: Scores for each group of strains as determined by pseudo-phylogeny and detected small molecules via IDBac. Groups were determined by cutting the dendrogram in **Figure S2A** at a height of 0.9. Scores (rounded to 2 decimal places) were then calculated for each strain using **Equation S1** based on the network shown. Strains selected for further analysis were selected and highlighted based on their scores and antiSMASH results as predicted using draft genomes.

| Group 1 | | | Group 2 | | |
|---|---|---|---|---|---|
| Strain | Score | Num of Predicted BGCs | Strain | Score | Num of Predicted BGCs |
| *G. arilaitensis* JB182 | 6.18 | 7 | *Brevibacterium* sp. EP18 | 2.48 | 7 |
| *E. coli* K12 | 3.04 | 2 | *Brevibacterium* sp. EP26 | 2.43 | 7 |
| *Hafnia alvei* JB232 | 1.15 | - | *Brachybacterium* sp. EP1 | 2.40 | 6 |
| *P. psychrophila* JB418 | 0.11 | 7 | *Brevibacterium linens* JB5 | 2.33 | 9 |
| | | | *Brevibacterium* sp. EP38 | 1.68 | 4 |
| | | | *Brevibacterium* sp. EP24 | 1.44 | 6 |
| | | | *Brevibacterium* sp. EP6 | 1.35 | 3 |
| | | | *Brachybacterium* sp. EP7 | 1.28 | 3 |
| | | | *Brevibacterium* sp. EP34 | 1.01 | 7 |
| | | | *Brevibacterium* sp. EP20 | 0.58 | 7 |
| | | | *Brevibacterium* sp. EP29 | 0.52 | 5 |
| | | | *Brevibacterium* sp. EP13 | 0.42 | 10 |
| | | | *Brevibacterium* sp. EP31 | 0.26 | 12 |
| | | | *Brevibacterium* sp. EP10 | 0.08 | 6 |
| | | | *Brachybacterium* sp. JB7 | 0.01 | 5 |

**Table S2**: Metabolites identified from fungal species and their level of identification as defined by Sumner *et al*.[7].

| Species | Compound Name | Molecular Formula | Calculated [M+H] | Measured [M+H] | ppm Error | PubChem CID | Level ID | Reference |
|---|---|---|---|---|---|---|---|---|
| Scopulariopsis sp. JB370 and 165-5 | Isariin A | $C_{33}H_{59}N_5O_7$ | 638.4493 | 638.4506 | 2.1 | 73424 | 4 | Saabareesh et al. 2007 |
| Scopulariopsis sp. JB370 and 165-5 | Isariin G2 | $C_{31}H_{56}N_5O_7$ | 610.4180 | 610.4195 | 2.5 | 44423145 | 3 | Saabareesh et al. 2007 |
| P. atramentosum | NBRI23477 A (Atpenin Analogue) | $C_{15}H_{21}Cl_2NO_5$ | 366.0875 | 366.0872 | 0.8 | 54741826 | 2 | Natural Products Atlas: NPA008775 |
| P. atramentosum | NBRI23477 B (Atpenin Analogue) | $C_{15}H_{21}NO_5$ | 296.1498 | 296.1498 | 0 | 5474182 | 3 | Natural Products Atlas: NPA009059 |
| P. atramentosum | Atpenin A4* | $C_{15}H_{23}ClNO_5$ | 322.1265 | 322.1269 | 1.2 | 54676867 | 2 | Natural Products Atlas: NPA008981 |
| P. atramentosum | Atpenin A5* | $C_{15}H_{21}Cl_2NO_5$ | 366.0875 | 366.0873 | 0.6 | 54676868 | 2 | Natural Products Atlas: NPA015919 |
| P. atramentosum | Atpenin B | $C_{15}H_{24}NO_5$ | 298.1655 | 298.1648 | 2.2 | 54676869 | 3 | Natural Products Atlas: NPA018005 |
| P. atramentosum | Glandicoline A | $C_{22}H_{21}N_5O_3$ | 404.1723 | 404.1732 | 2.3 | 91820007 | 3 | |
| P. atramentosum | Glandicoline B | $C_{22}H_{21}N_5O_4$ | 420.1672 | 420.1656 | 3.7 | 124079399 | 3 | |
| P. atramentosum | Roquefortine C | $C_{22}H_{23}N_5O_2$ | 390.1930 | 390.1940 | 2.6 | 21608802 | 3 | |
| P. atramentosum | Andrastatin A | $C_{28}H_{38}O_7$ | 487.2696 | 487.2684 | 2.4 | 6712564 | 2 | GNPS: CCMSLIB00004713231 CCMSLIB00004713240 |
| P. atramentosum | Citreohybridonol | $C_{28}H_{36}O_8$ | 501.2489 | 501.2486 | 0.4 | 101252260 | 2 | GNPS: CCMSLIB00004711830 |
| P. atramentosum | Meleagrin | $C_{23}H_{23}N_5O_4$ | 434.1828 | 434.1827 | 0.3 | 23728435 | 2 | GNPS: |

| | | | | | | | | |
|---|---|---|---|---|---|---|---|---|
| | | | | | | | | CCMSLIB00000852096 |
| P. atramentosum | Oxaline | $C_{24}H_{25}N_5O_4$ | 448.1985 | 448.1982 | 0.6 | 70698220 | 2 | MassBank EU: AC000745 |
| P. solitum | Atlantinone A | $C_{26}H_{34}O_6$ | 443.2434 | 443.2436 | 0.5 | 101516467 | 2 | GNPS: CCMSLIB00000853278 |
| P. solitum | Cyclopenol | $C_{17}H_{14}N_2O_4$ | 311.1032 | 311.1021 | 3.5 | 101201 | 2 | GNPS: CCMSLIB00000852686 |
| P. solitum | Viridicatol | $C_{15}H_{11}NO_3$ | 254.0817 | 254.0819 | 0.7 | 115033 | 2 | GNPS: CCMSLIB00000845922 |
| P. solitum | Pyripyropene O (Putative Adduct) | $C_{29}H_{35}NO_7$ | $[M+NH_4]$ = 527.2757 | $[M+NH_4]$ = 527.2861 | 18 | 10553713 | 3 | GNPS: CCMSLIB00000848971 |
| P. solitum | Pyripyropene Analog | $C_{29}H_{35}NO_6$ | $[M+NH_4]$ = 511.2808 | $[M+NH_4]$ = 511.2906 | 19 | | 3 | GNPS: CCMSLIB00000848971 |
| P. solitum | ML-236A | $C_{18}H_{26}O_4$ | 307.1909 | 307.1901 | 2.6 | 173651 | 2 | GNPS: CCMSLIB00000478442 |
| P. solitum | Cyclopeptine | $C_{17}H_{16}N_2O_2$ | 281.1290 | 281.1284 | 2.1 | 15649435 | 2 | GNPS: CCMSLIB00000854845 |
| P. atramentosum, P. camemberti, P. solitum | Desferrichrome | $C_{27}H_{45}N_9O_{12}$ | 688.3266 | 688.3234 | 4.6 | 169636 | 1 | GNPS: CCMSLIB00005435755 |
| P. atramentosum, P. camemberti, P. solitum | Desferricoprogen | $C_{35}H_{56}N_6O_{13}$ | 769.3984 | 769.3976 | 1 | 23636677 | 1 | |
| F. domesticum | Desferricoprogen B | $C_{33}H_{54}N_6O_{12}$ | 727.3878 | 727.3882 | 0.5 | 122198275 | 2 | GNPS: CCMSLIB00004679227 |
| F. domesticum | Palmitoylcoprogen | $C_{49}O_{13}H_{82}N_6Fe$ | 1018.5289 | 1018.5243 | 4.5 | | 3 | |

**Table S3**: Number of genes in species specific gene sets with enriched functions in pairwise co-cultures of native cheese rind fungi and *P. psychrophila* JB418 as reported by Pierce *et al*.[6].

| Fungal Growth Partner | Number of Genes |
|---|---|
| *Scopulariopsis* sp. JB370 | 85 |
| *Scopulariopsis* sp. 165-5 | 269 |
| *Penicillium atramentosum* RS17 | 328 |
| *Penicillium camemberti* SAM3 | 516 |
| *Penicillium solitum* #12 | 463 |
| *Fusarium domesticum* 554A | 56 |
| *Diutina catenulata* 135E | 80 |
| *Debaromyces hansenii* 135B | 34 |

**Table S4**: Number of genes in species specific gene sets with enriched functions in pairwise co-cultures of native cheese rind fungi and *E. coli* K12 as reported by Pierce *et al*.[6].

| Fungal Growth Partner | Number of Genes |
|---|---|
| *Scopulariopsis* sp. JB370 | 260 |
| *Scopulariopsis* sp. 165-5 | 130 |
| *Penicillium atramentosum* RS17 | - |
| *Penicillium camemberti* SAM3 | 12 |
| *Penicillium solitum* #12 | 51 |
| *Fusarium domesticum* 554A | 220 |
| *Diutina catenulata* 135E | 185 |
| *Debaromyces hansenii* 135B | 124 |

**Table S5**: Metabolites identified from *Actinobacteria* species and their level of identification as defined by Sumner *et al*.[7]. Note that ppm error is only reported for library matches that were not determined to be analogues. Otherwise, the mass differences are reported instead. Although the timsTOF fleX is capable of reaching four significant features, only three have been reported due to being the default in GNPS. The GNPS job can be found at https://gnps.ucsd.edu/ProteoSAFe/status.jsp?task=570905801277433a88add3bb735c963c.

| Species | Compound Name | Molecular Formula | Calculated [M+H] | Measured [M+H] | ppm Error | Mass Difference | Pubchem CID | Level ID | Reference |
|---|---|---|---|---|---|---|---|---|---|
| Arthonia endlicheri | Arthonin Analogue | $C_{25}H_{32}N_2O_4$ | Arthonin = 425.244 | 340.166 | | 85.078 | 11101959 | 3 | Natural Products Atlas: NPA009730 |
| Aspergillus versicolor | Cyclo-(D)-Tyr-(D)-Pro Analogue | $C_{14}H_{16}N_2O_3$ | Cyclo-(D)-Tyr-(D)-Pro = 261.124 | 327.135 | | 66.011 | 28125534 | 3 | Natural Products Atlas: NPA000159 |
| Streptomyces sp. KH29 | Cyclo-(L)-Trp-(L)-Trp Analogue | $C_{22}H_{20}N_4O_2$ | Cyclo-(L)-Trp-(L)-Trp = 373.166 | 286.155 | | 87.011 | 7091706 | 3 | Natural Products Atlas: NPA002698 |
| Streptomyces sp. KH29 | Cyclo-(L)-Trp-(L)-Trp Analogue | $C_{22}H_{20}N_4O_2$ | Cyclo-(L)-Trp-(L)-Trp = 373.166 | 284.14 | | 89.026 | 7091706 | 3 | Natural Products Atlas: NPA002698 |
| Fungus | Laxaphycin D | $C_{63}H_{110}N_{14}O_{19}$ | $[M+2H]^{2+}$ = 684.414 | $[M+2H]^{2+}$ = 684.393 | 30.7 | 0.021 | 101632355 | 3 | Natural Products Atlas: NPA014810 |
| Sponge | Makaluvamine I Analogue | | Makaluvamine I = 188.080 | 226.098 | | 38.018 | 135474607 | 3 | GNPS: CCMSLIB00004679175 |
| Bacillus pumilus | N-Acetyltryptophan Analogue | $C_{13}H_{14}N_2O_3$ | N-Acetyltryptophan = 247.108 | 323.139 | | 76.031 | 2002 | 3 | Natural Products Atlas: NPA015473 |

| | | | | | | | | | |
|---|---|---|---|---|---|---|---|---|---|
| Bacillus pumilus | N-Acetyltryptophan Analogue | $C_{13}H_{14}N_2O_3$ | N-Acetyltryptophan = 247.108 | 339.134 | | 92.026 | 2002 | 3 | Natural Products Atlas: NPA015473 |
| Xenorhabdus nematophilus | Nematophin Analogue | $C_{16}H_{20}N_2O_2$ | Nematophin = 273.160 | 189.103 | | 84.057 | 9881952 | 3 | Natural Products Atlas: NPA007705 |
| Xenorhabdus sp. PB62.7 | Nevaltophin A Analogue | | Nevaltophin A = 347.230 | 383.176 | | 35.946 | 132509308 | 3 | GNPS: CCMSLIB00000840595 |
| Xenorhabdus sp. PB62.7 | Nevaltophin D Analogue | $C_{22}H_{31}N_3O_3$ | Nevaltophin D = 386.244 | 422.187 | | 35.943 | 132509319 | 3 | Natural Products Atlas: NPA0028349 and GNPS: CCMSLIB00000840594 |
| Candida albicans | Tryptophol Analogue | $C_{10}H_{11}NO$ | Tryptophol = 162.092 | 203.118 | | 41.026 | 10685 | 3 | Natural Products Atlas: NPA006412 |
| Candida albicans | Tryptophol Analogue | $C_{10}H_{11}NO$ | Tryptophol = 162.092 | 261.124 | | 99.032 | 10685 | 3 | Natural Products Atlas: NPA006412 |
| Rhizopogon roseolus | Zeatin Riboside | $C_{15}H_{21}N_5O_5$ | 352.162 | 352.162 | 0 | 0 | 6440982 | 2 | Natural Products Atlas: NPA009273 |
| Rhizopogon roseolus | Zeatin Riboside Analogue | $C_{15}H_{21}N_5O_5$ | Zeatin Riboside = 352.162 | 398.15 | | 45.988 | 6440982 | 3 | Natural Products Atlas: NPA009273 |
